# A deep profile of gene expression across 18 human cancers

**DOI:** 10.1101/2024.03.17.585426

**Authors:** Wei Qiu, Ayse B. Dincer, Joseph D. Janizek, Safiye Celik, Mikael Pittet, Kamila Naxerova, Su-In Lee

## Abstract

Clinically and biologically valuable information may reside untapped in large cancer gene expression data sets. Deep unsupervised learning has the potential to extract this information with unprecedented efficacy but has thus far been hampered by a lack of biological interpretability and robustness. Here, we present DeepProfile, a comprehensive framework that addresses current challenges in applying unsupervised deep learning to gene expression profiles. We use DeepProfile to learn low-dimensional latent spaces for 18 human cancers from 50,211 transcriptomes. DeepProfile outperforms existing dimensionality reduction methods with respect to biological interpretability. Using DeepProfile interpretability methods, we show that genes that are universally important in defining the latent spaces across all cancer types control immune cell activation, while cancer type-specific genes and pathways define molecular disease subtypes. By linking DeepProfile latent variables to secondary tumor characteristics, we discover that tumor mutation burden is closely associated with the expression of cell cycle-related genes. DNA mismatch repair and MHC class II antigen presentation pathway expression, on the other hand, are consistently associated with patient survival. We validate these results through Kaplan-Meier analyses and nominate tumor-associated macrophages as an important source of survival-correlated MHC class II transcripts. Our results illustrate the power of unsupervised deep learning for discovery of cancer biology from existing gene expression data.

## Introduction

Gene expression profiles are the reflections of a complex network of underlying cellular and molecular processes. Unsupervised learning is a key step toward extracting meaningful biological information from expression profiles and reducing the dimensionality of the data for downstream tasks, such as prediction of phenotypes. Unsupervised learning projects high-dimensional input variables into a latent space consisting of a smaller set of latent variables, or factors, capable of explaining the variation in the original input space. Learned latent variables represent sources of genome-wide expression variation across samples, for example large-scale transcriptional programs that define intrinsic disease subtypes or reflect extrinsic stimuli such as hypoxia or treatment pressure. Each individual cancer has different characteristics and response to therapy, even cancers of the same type. Therefore, discovering and understanding biologically meaningful sources of expression variation is of considerable interest from a research and clinical perspective.

One key limitation of commonly used latent space learning approaches for expression data, such as principal component analysis (PCA), is that they can only extract latent variables that have linear relationships with gene expression levels, while gene interactions can be more complex. The artificial intelligence (AI) field has achieved notable success in unsupervised learning by using deep neural networks that can capture highly complex relationships between variables. It has been shown that the latent variables extracted by unsupervised deep learning approaches from image data represent high-level features that are intuitively important for the entire image in the training set, for example: skin color, age, and gender from face images^1^, lighting and room geometry from scene images^2^, and rotation and size of an object from 3D images^3^. These informative and complex image features cannot be captured by models limited to learning linear feature interactions^4^.

The success of unsupervised deep learning in computer vision has motivated several recent applications of deep unsupervised learning methods to gene expression profiles. Prior approaches have used generative modeling to learn the latent factors underlying single cell sequencing data, separating technical artifacts from biological factors^5^. Furthermore, previous studies have conducted pan-cancer analyses with various approaches ranging from co-expression networks, to differential expression analysis, to deep unsupervised learning approaches^6–15^. For example, Kim et al. (2020) introduced a deep learning architecture to enable transfer learning of unsupervised deep models to improve survival prediction and applied it to The Cancer Genome Atlas (TCGA) data. Way & Greene (2018) pioneered the application of unsupervised deep learning to capture biologically relevant features from TCGA expression data.

However, three challenges still impede the successful application of deep unsupervised learning approaches to cancer expression data. First, deep learning has a high risk of overfitting when not provided with large sample numbers. Second, the non-deterministic nature of the learning process impairs the robustness of the learned latent spaces. Each run of neural network training, even using the same architecture, results in different models with different parameters, which makes it difficult to capture consistent signals^16^. Model consistency is of paramount importance in biology, where interpretation of the learned model is more important than obtaining high predictive accuracy. Third, neural networks with multiple hidden layers are “black boxes” by nature: since it is not clear how the model uses gene expression inputs to generate a latent variable, biological interpretation of latent variables is problematic.

In addressing the inherent non-determinism in training deep learning models, particularly for biological data analysis, model ensembles emerge as a potent solution. By aggregating outputs from multiple model runs, ensembles enhance the consistency and stability of predictions, crucial for biological applications^17–19^. Whereas prior techniques have suggested the use of model ensembles in unsupervised learning^16,20^, these methods have so far been limited to “shallow” models with a single hidden layer. Moreover, the application of Explainable AI (XAI) in life sciences^21–23^, although widespread, often grapples with complex, multidimensional data. In this context, model ensembles offer a substantial advantage, improving the quality and reliability of feature attribution^24^, thereby aligning with the growing emphasis on transparency and comprehension in AI models used for biological data analysis.

To resolve these challenges, we developed DeepProfile, a framework that enables a unique pan-cancer analysis by learning statistically robust and interpretable latent spaces from gene expression data (**Fig. 1**). To robustly train the neural networks, we incorporated expression datasets comprising 18 human cancers from 50,211 transcriptomes in the public gene expression data repository Gene Expression Omnibus (GEO)^25^. To address the non-deterministic nature of the deep learning process and capture robust latent spaces, we devised a unique ensemble approach to integrate the results of hundreds of deep unsupervised models generated from different random starting points and latent space sizes. While previous approaches have proposed using ensembles of models^16,20^, these methods have so far been limited to “shallow” unsupervised models with a single hidden layer. By incorporating state-of-the-art feature attribution methods that can provide gene importance values for each latent variable, DeepProfile is able to create ensembles of “deep” unsupervised models with multiple hidden layers. Finally, DeepProfile extends previous studies by incorporating an extended set of gene expression profiles from GEO, The Cancer Genome Atlas (TCGA)^26^, and the Genotype-Tissue Expression (GTEx) database^27^, and by integrating different data modalities such as clinical and mutational features. This rich resource of robust cancer-specific deep embeddings, the values of the latent variables, and biological characterization of the latent variables enables us to examine cancer transcriptomic signals from a new angle and investigate their associations to various phenotypes.

**Fig. 1.**
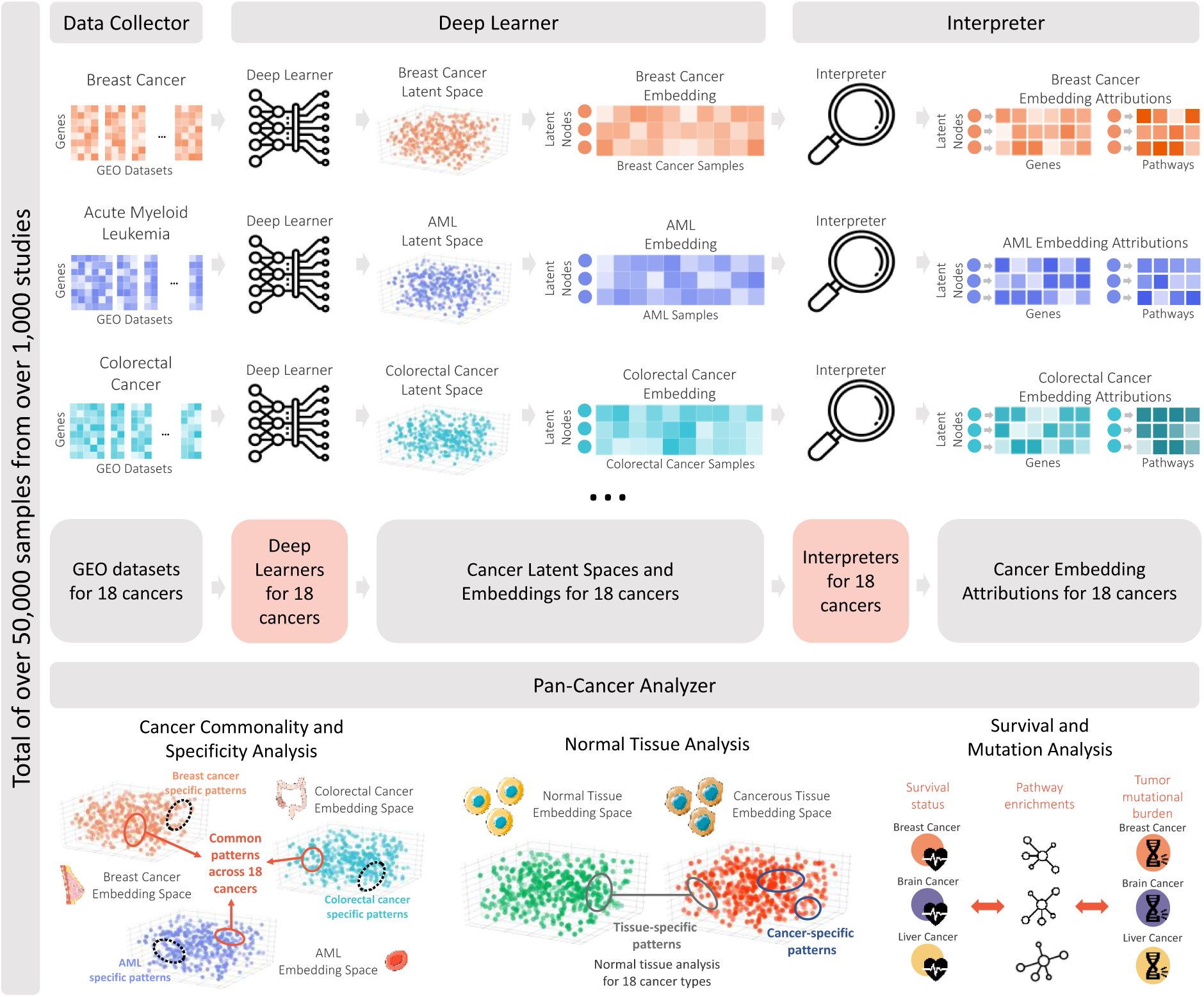
DeepProfile Pan-Cancer Framework. **Data Collector:** We downloaded gene expression datasets for 18 cancer types from the common microarray platforms, preprocessing and concatenating them into cancer-specific expression matrices. In total, we have over 50,000 samples from over 1,000 GEO datasets. **Deep Learner:** We pass the expression matrices to Deep Learner models to learn cancer-specific latent spaces. Deep Learner is an ensemble of variational autoencoders (VAEs) that encodes the high-dimensional expression signals to a biologically informative *latent space*. We then map the training samples to the learned latent spaces and define cancer sample *embeddings*, where each DeepProfile latent variable encodes a certain source of variance across cancer samples. **Interpreter:** We pass the learned embeddings to Interpreter models to extract *gene-level and pathway-level attributions* for each latent variable. Gene-level attributions denote how much each gene contributes to a latent variable. Similarly, pathway-level attributions denote the pathways significantly associated with the most important genes of each latent variable. **Pan-Cancer Analyzer:** Using the cancer-specific embeddings and attributions; we carry a detailed pan-cancer analysis including **(1)** analyzing the latent spaces of 18 cancers to discover cancer-common and specific patterns, **(2)** differentiating cancer-specific patterns from tissue-specifying ones by contrasting cancer embeddings to normal tissue embeddings and **(3)** investigating survival and mutation related signals by integrating DeepProfile embeddings with survival and tumor mutational burden profiles **(See Extended Data Fig. 1 and 2)**.

Using the DeepProfile framework, we examine genes and pathways that capture major variation across all 18 cancer types. We find that universally important genes control aspects of the inflammatory response by modulating the transcriptional phenotypes of tumor-infiltrating immune cells. Cancer-type specific genes with large contributions to the latent spaces of only one particular cancer type, on the other hand, define molecular disease subtypes and reflect tissue-specific biology. We develop a methodology for linking DeepProfile embeddings to patient- and tumor-level characteristics and apply the method to study genes and pathways that – as seen through the lens of DeepProfile’s latent spaces – correlate with tumor mutation burden and patient survival. We find that tumor mutation burden is significantly associated with the expression of cell cycle-related pathways across a large majority of cancers, while survival correlates with DNA mismatch repair and MHC class II antigen presentation pathway activity. Our methodologies to make deep neural network models biologically interpretable allow for complex, non-linear relationships to be learned while retaining stable models. Thus, DeepProfile’s robustness and interpretability enables the discovery of unique biological patterns in large gene expression datasets.

## Results

### DeepProfile learns robust latent spaces for 18 cancer types

Because highly expressive models such as deep neural networks tend to overfit when the sample size is small, we obtained all available expression datasets from the most common microarray platforms for 18 human cancers from GEO^25^ (**Fig. 1; Supplemental File 1**) (**see Methods)**, resulting in 50,211 samples from 1,098 datasets. DeepProfile projects the expression data into lower-dimensional latent space represented by a set of latent variables using an ensemble approach for the variational autoencoder (VAE)^28^ (**Extended Data Fig. 1**). The VAE is a special type of deep neural network that compresses high-dimensional data (here, tens of thousands of genes) into low-dimensional embeddings with minimal information loss. More specifically, two neural networks – (i) the encoder that models the relationship between input variables and latent variables in the latent space and (ii) the decoder that models the relationship between the latent variables and the reconstructed input variables – are trained such that the reconstructed input data are close to the input gene expression data (**see Methods**).

VAE is a unique model that can discover non-linear relations among genes to reflect the true nature of gene interactions. However, applying the model to expression data is not straightforward. Neural networks inherently suffer from learned model variability across different random initializations due to their intrinsic non-convex nature. This means that a conventional learning algorithm for VAE can result in a model that is different in every trial, an outcome that hinders the inference of robust biological signals. To improve robustness, we developed an ensemble of VAEs to combine the learned models from different random runs and latent dimension sizes (**Extended Data Fig. 1**) (**see Methods**). This approach integrates signals from hundreds of different latent spaces into one information-rich space. After learning these cancer-specific latent spaces, DeepProfile’s ‘interpreter’ biologically characterizes each latent variable by mapping it to genes and pathways (**Fig. 1**). This process is based on the principled ‘feature attribution’ method, namely integrated gradients^29^, to quantify how much each latent variable’s value is attributed to input variables (**Fig. 1** and **Extended Data Fig. 2**). In particular, for each latent variable, DeepProfile produces a list of gene attribution scores, which indicate the relevance of each gene to that latent variable and uses the top-listed genes for pathway enrichment tests, which provide pathway-level attribution scores (**see Methods**).

The input gene expression datasets, their lower-dimensional embeddings, gene-level and pathway-level relevance, and the results of our pan-cancer analysis are publicly available at: https://github.com/suinleelab/deepprofile-study (code), and https://doi.org/10.6084/m9.figshare.25414765.v2 (data).

The trained DeepProfile model explains the relevant factors of gene expression variation in each sample by encoding high-dimensional measurements of thousands of gene expression levels into 150 latent variables. The number of latent variables was determined using an algorithm that iteratively decides whether to add an additional latent variable using a statistical test of Gaussianity (see **Methods**). DeepProfile can be applied to any new cancer gene expression dataset to reduce its dimensionality (**Extended Data Fig. 2; Methods**). To demonstrate the consistency with independent RNA-Seq data, we used RNA-seq data from TCGA^26^ containing 9,079 samples across 18 cancers which were not used for training DeepProfile (**Extended Data Fig. 2**) (**see Methods**; **Supplementary File 1**). Our result also highlights that DeepProfile can be successfully applied to RNA-Seq expression profiles despite being trained on microarray data. This is further supported by the high correlation between DeepProfile embeddings generated from microarray and RNA-seq data (**Extended Data Fig. 3**).

### DeepProfile can learn biologically interpretable latent variables enriched for a wide set of pathways

It is desirable for latent variables to be biologically interpretable. DeepProfile provides gene attribution scores for each latent variable, thereby enabling a standard enrichment test to assess the overlap’s statistical significance using the Fisher’s exact test between the top-scoring genes and predefined gene sets, available through curated pathway databases such as KEGG, BioCarta, and Reactome. Pathway annotation dramatically facilitates the interpretation of a latent variable’s biological meaning; ideally, latent variables will capture many known pathways. We compared the average number of pathways captured by DeepProfile’s latent variables to results from other dimensionality reduction methods (**see Methods**). DeepProfile latent variables captured more pathways than alternative methods (106 test cases out of 108, proportions z-test P = 1.62 × 10^−301^) (**Fig. 2a** top). Further, when we focused on oncogenic pathways (as defined by MSigDb) specifically, DeepProfile outperformed the other methods in terms of total gene sets captured (102 tests cases out of 108, proportions z-test P = 2.03 × 10^−90^) (**Fig. 2a** bottom). This means DeepProfile not only captures more pathways but also identifies the pathways relevant to cancer. **Supplementary Fig. 1** illustrates that 156 pathways are significant in more than nine cancer types, highlighting a significant overlap of common pathways across cancers. Additionally, we identified 163 pathways unique to individual cancer types, emphasizing the specificity of the DeepProfile approach in detecting cancer-specific biological signals. We also evaluated the uniqueness and redundancy of pathways identified by DeepProfile’s latent variables in **Supplementary Note 1**, revealing the model’s proficiency in distinguishing unique biological variations.

**Fig. 2.**
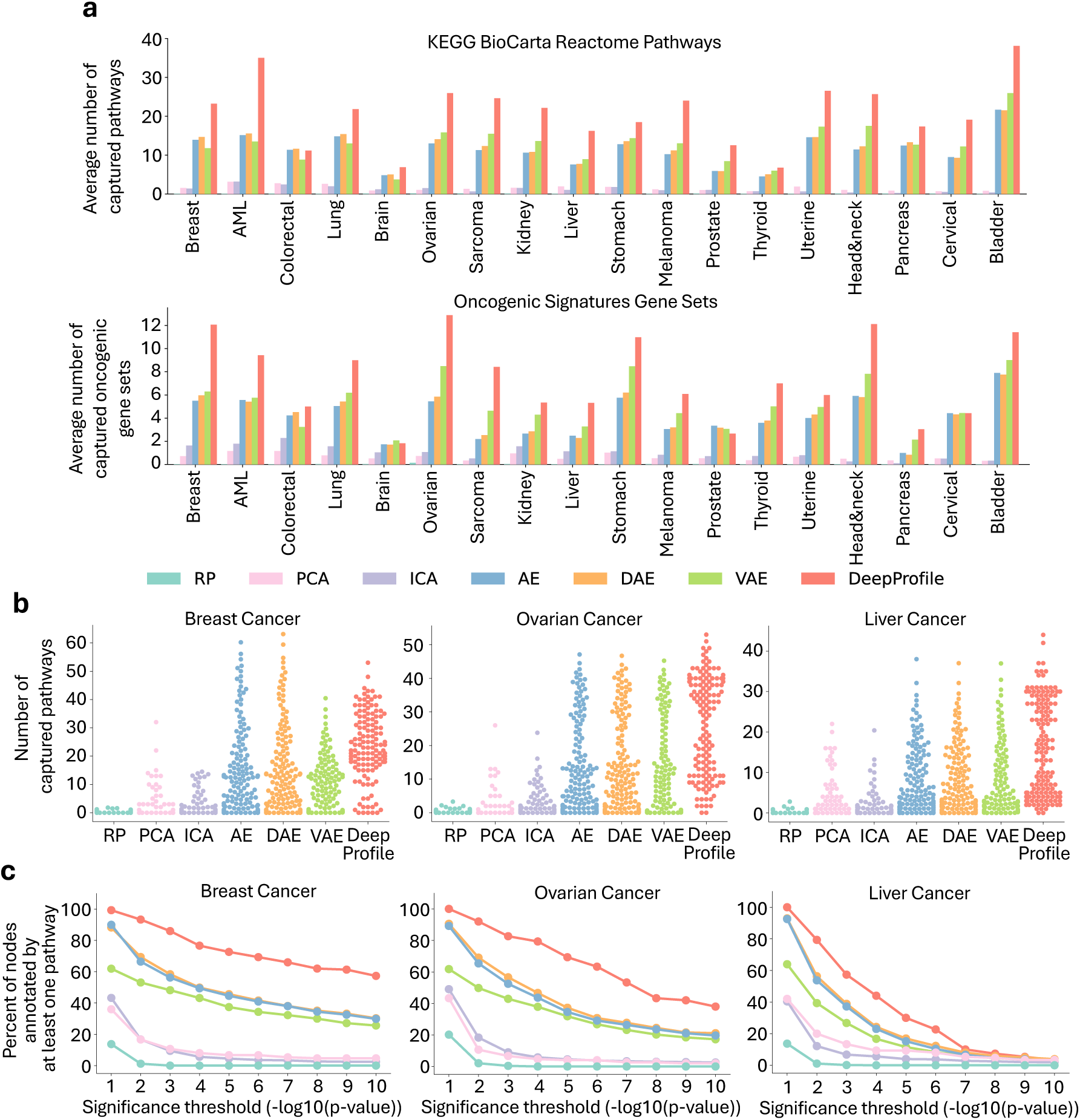
Comparison of pathway enrichment of DeepProfile and other dimensionality reduction methods. **a.** The average number of pathways significantly captured (FDR corrected p-value < 0.05) by latent variables of latent embeddings of DeepProfile and other dimensionality reduction methods are shown for KEGG, BioCarta, Reactome pathways **(Top Plot)** and Oncogenic Signatures gene sets **(Bottom Plot)**. Each latent variable of each embedding is associated with each pathway with a p-value and we count the number of pathways significantly captured by each latent variable. We then average these pathway counts over all latent variables to define the average number of pathways significantly captured by a method. **b.** Distribution plots of number of KEGG, BioCarta, Reactome pathways significantly captured (FDR corrected p-value < 0.05) by each latent variable shown for 3 cancer types (**see Extended Data Fig. 4** for all 18 cancers). **c.** Comparison of the percent of latent variables annotated by at least one pathway above the significance threshold. The percent of annotated latent variables are shown for multiple significance thresholds for DeepProfile and alternative dimensionality reduction methods. Examples from 3 cancer types are provided (**see Extended Data Fig. 5** for all 18 cancers).

A latent variable not associated with any known pathway is difficult to characterize biologically, thus decreasing overall interpretability. We found that DeepProfile produces fewer such pathways than other methods (**Fig. 2b** and **Extended Data Fig. 4**) (**see Methods**). Further, we show that, for varying p-value thresholds, a higher percentage of DeepProfile latent variables are biologically annotated compared to other methods **(Fig. 2c** and **Extended Data Fig. 5**) (**see Methods**). To validate DeepProfile’s discriminatory power against random patterns, we explored its performance on Gaussian noise datasets, simulating conditions devoid of actual biological signals. The results highlight the model’s precision in differentiating genuine biological signals from noise (**Supplementary Note 2**). These results demonstrate that DeepProfile’s unique deep learning ensemble approach improves latent variables’ biological interpretability. Using the robustly identified latent space and embeddings and the gene-level and pathway-level interpretation of each latent variable, we next proceeded to perform in-depth analyses of the biology revealed by DeepProfile.

### Universally important genes modulate inflammatory pathways

We began by investigating genes with universally large gene attribution scores to DeepProfile latent variables across all cancer types (**see Methods, Supplementary Files 2** and **3**). These genes represent dominant gene expression programs that consistently explain considerable portions of the transcriptional variance across many different cancers. We found that universal genes with high average gene attribution scores were primarily involved in immune response regulation and antigen presentation (35 out of the top universal 100 genes, p-value: 9.4 × 10^−6^) (**Figs. 3a-c**). Given that solid tumors (which constitute most of our data) can be infiltrated by immune cells to varying degrees, we hypothesized that universal genes may reflect the gene expression signatures of various admixing immune cell types. To test this hypothesis, we assessed the overlap between four signatures of major immune cell types (T cells, B cells, neutrophils, and macrophages; **see Methods, Supplementary File 4**)^30^ and genes with top DeepProfile attribution scores (**see Methods**). We found that there was a small overlap between top DeepProfile genes and the macrophage signature (2 out of the top universal 100 genes, p-value: 2.5 × 10^−2^), but not any of the other immune cell type signatures. To enhance our analysis, we utilized pre-computed immune cell fractions from the TCGA data^31^. We calculated Pearson’s correlations between gene expression levels and the proportions of various immune cell types, identifying the top 100 genes most correlated with each cell type. Subsequent Fisher’s exact tests showed minimal or no significant overlap between these correlation-based top genes and the top 100 DeepProfile genes (**Supplementary Table 1**, **Methods**), suggesting that the gene sets driving immune infiltration are distinct from those identified by DeepProfile’s signatures.

**Fig. 3.**
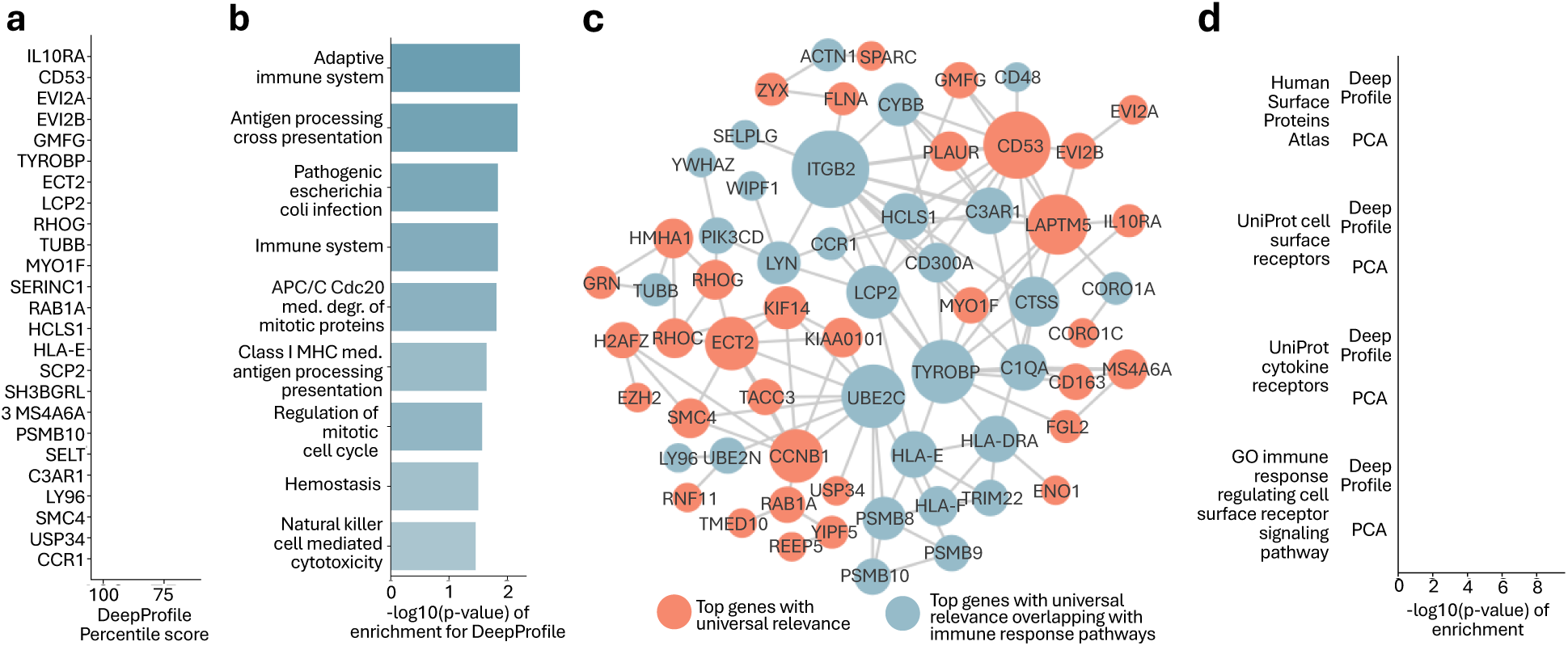
DeepProfile cancer commonality analysis. **a.** List of top highest-scoring genes across 18 cancer types for DeepProfile. The percentile scores of the top scoring genes are shown for all cancers and the average percentile scores across 18 cancers are highlighted. We calculated the average importance of a gene for DeepProfile embedding by calculating the average gene importance scores across all latent variables of the embedding, converting the average importance scores to percentile scores and averaging these percentile scores across all 18 cancers. The plot is zoomed in for clear comparison. The universal importance scores for all genes are available in **Supplementary File 2**. **b.** The top enriched pathways (KEGG, BioCarta, Reactome) for the top 100 universally important DeepProfile genes and the corresponding FDR-corrected p-values. Enrichment scores for all pathways are available in **Supplementary File 2**. **c.** Network of top 100 genes with universal importance. The network is generated with StringDB and disconnected latent variables are excluded. The size of a latent variable is determined by hubness, i.e., the number of edges. Genes that are included in immune response related pathways are colored blue. **d.** The enrichment p-values for cell surface and cytokine receptors for DeepProfile and PCA top 100 universally important genes.

Next, we hypothesized that DeepProfile prioritized genes whose expression was associated with recurrent transcriptional phenotypes in tumor-infiltrating immune cells, such as signatures linked with immune cell activation or suppression. To illustrate this concept, consider the gene with the highest average attribution, the alpha subunit of the interleukin 10 receptor (*IL10RA*). *IL10RA* scored among the top 1% of genes in 14 out of 18 cancers (78% of cancer types) and top 10% in all 18 cancer types, indicating that DeepProfile consistently ascribed high explanatory power to this gene, regardless of tissue context (**Fig. 3a**). Upon encountering an inflammatory stimulus, a variety of immune cells upregulate *IL10RA*, which mediates the activation of a compensatory anti-inflammatory gene expression program; *IL10RA* has consequently been described as a “master switch” regulating the balance between pro- and anti-tumor inflammation^32^. Therefore, transcript levels of IL10RA do not only reflect the presence or absence of IL10RA-expressing immune cells, they also predict several thousand genes regulated by IL10RA^33^, potentially explaining the large role this gene plays in DeepProfile’s latent spaces.

To test the hypothesis that universally high-scoring DeepProfile genes were enriched for transcripts that, like IL10RA, modulate immune cells’ transcriptional phenotypes, we quantified cell surface receptors among genes with top attribution scores. We reasoned that cell surface receptors are enriched for proteins that relay extra-cellular signals and thus have the potential to regulate immune cells’ transcriptional phenotypes. We collected gene sets containing cell surface proteins and receptors from the Cell Surface Protein Atlas (CSPA)^34^, the UniProt database^35^, and the Gene Ontology database (GO)^36^ (**Supplementary File 5**). We found highly significant overlap between these gene sets and genes with top average DeepProfile attribution scores across all cancers (15, 32, and 12 out of the top universal 100 genes, respectively; p-values: 1.5 × 10^−5^, 7.0 × 10^−10^, 1.0 × 10^−5^) (**Fig. 3d**) (see **Methods**). Importantly, PCA did not recover these cell surface proteins and receptors (**Fig. 3d; Supplementary File 2**) (**see Methods**), indicating that DeepProfile’s ability to identify non-linear relationships is essential in capturing this source of variance, and that the functional relationship between receptor expression and gene expression modulation may itself have non-linear form.

In addition to IL10RA, DeepProfile’s top attributions contained many lesser known but potentially important genes that are consistently involved in the latent spaces of most cancer types. These included CD53, an immune-cell specific tetraspanin^37^; EVI2A and EVI2B, genes that control granulocytic differentiation^38^; and TYROBP, an adaptor protein that in association with various receptors mediates immune cell activation^39^ (**Fig. 3a**). As indicated above, none of these genes appear to signal the presence of a particular immune cell type in the tumor microenvironment, as they are broadly expressed by many different cells, but instead may be involved in modulating tumor-resident immune cells’ transcriptional phenotypes.

### Universally important pathways include cell cycle, immune system, and oxidative phosphorylation

Next, to investigate pathway-level information captured by DeepProfile, we studied the relationship between the embeddings and curated pathway gene sets available through the KEGG, BioCarta, and Reactome databases (see **Supplementary File 3**). We considered a pathway to be significantly enriched in a given cancer type if it overlapped with an FDR-corrected p-value below 0.05 with at least one DeepProfile latent variable (see **Methods**). We then extracted the pathways captured in the largest number of cancer types, grouped these pathways by functional category, and sorted the categories by the average number of cancer types in which they were significantly detected.

As expected, cell cycle-related gene sets were almost universally important, confirming that differences in proliferative index are a major source of variation across cancer transcriptomes (**Fig. 4**). This observation is consistent with long-standing clinical experience - some cancers evidently have higher mitotic rates than others - and the cell cycle consequently is found to play a role in nearly every morphological or molecular characterization of cancer^40–44^. Two cancer types had notably less pronounced contributions from cell cycle-related gene sets: AML, whose latent space mainly captured pathways related to adaptive immune response, and thyroid cancer, for which the most important pathways were related to mitochondrial function (**Supplementary File 3**). The two most common types of thyroid cancer (papillary and follicular) are exceptionally slow-growing neoplasms, which may explain this relative lack of contribution by cell cycle-related pathways. In AML, growth rates are more difficult to assess^45^, but it may be that most patients experience uniformly high growth rates due to the disease’s aggressiveness and its lack of spatial restraint. In both cases, a lack of variation in proliferative fractions across patients would explain why DeepProfile did not detect the cell cycle as an important contributor of variance in these cancers’ transcriptomes.

**Fig. 4.**
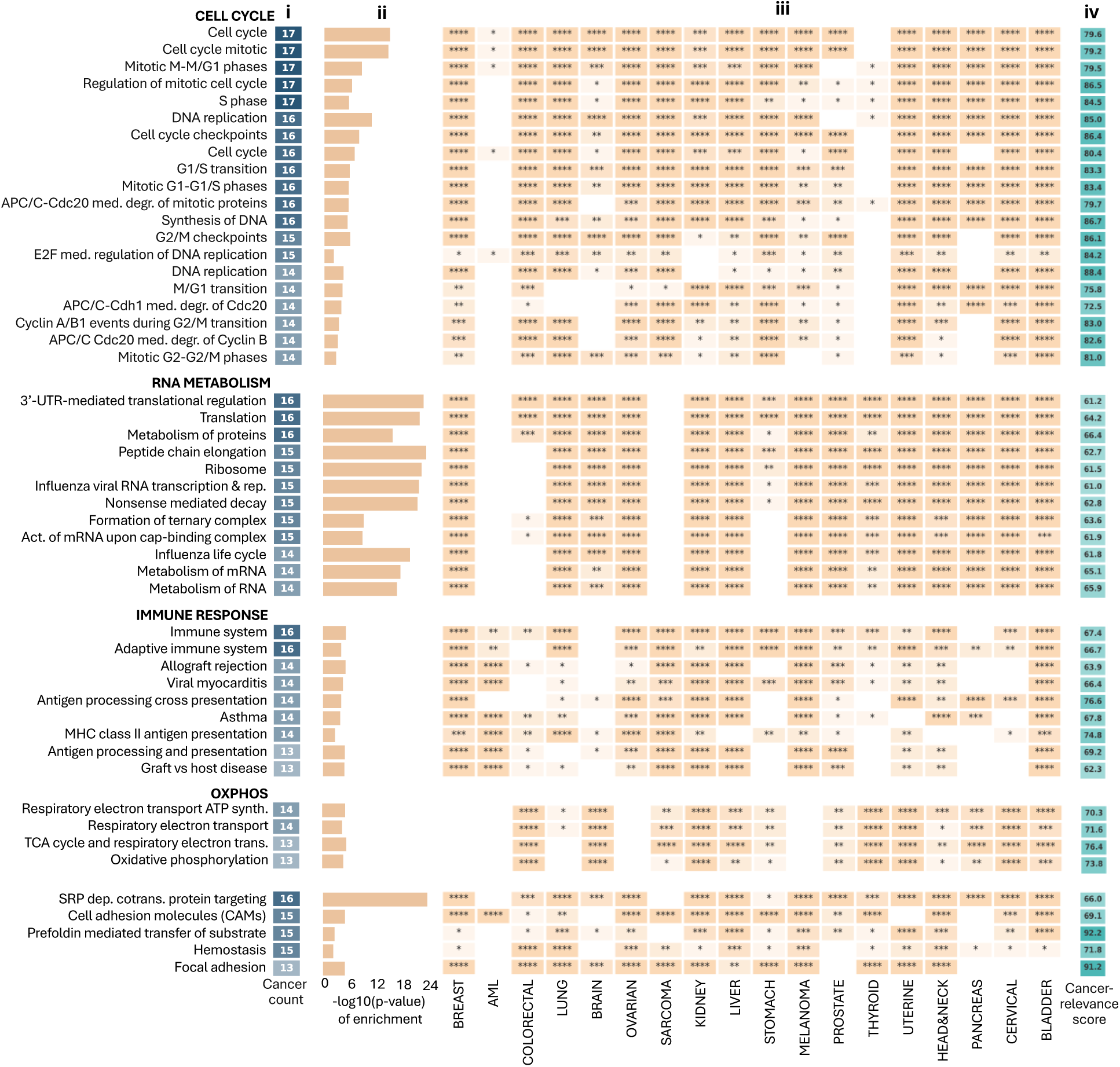
List of top KEGG, BioCarta, Reactome pathways that are universally important. The pathways are sorted based on the number of cancer types significantly capturing the pathway. All the scores for all pathways are available in **Supplementary File 3**. **i** Number of cancer types (out of 18) significantly capturing (FDR-corrected p-value < 0.05) each pathway. **ii** –log10(p-value of enrichment) averaged over all cancers significantly capturing the pathway. **iii** Heatmap denoting the significance of enrichment p-values for top pathways and all cancer types. The star annotations correspond to the significance of enrichment (* = p-value < 0.05, ** = p-value < 0.01, *** = p-value < 0.001, **** = p-value < 0.0001). **iv** Cancer character scores of pathways. The cancer character score denotes the relevance of each pathway to normal or cancerous tissue where a higher score indicates that the pathway is specifically important for cancerous tissues. The pathways are grouped manually in terms of their functional relations. The order of the groups is determined by the average cancer character score of each pathway group.

Immune-related pathways, as discussed in detail above, were the third-most frequently captured category (**Fig. 4**) followed by gene sets related to oxidative phosphorylation (OXPHOS), indicating that individual tumors’ position on the metabolic continuum between glycolysis and aerobic respiration explains global differences in their gene expression profiles^46^. Genes related to RNA metabolism and ribosome function also emerged as relevant across a large number of cancers; enrichment p-values were particularly significant in this category (**Fig. 4**). Consistent with prior pan-cancer analyses^11,43,44,47^, our study reinforces the significance of both immune-related and metabolism-related pathways across various cancer types, underlining their critical role in cancer biology. The identification of these well-established pathways initially validates the effectiveness of our approach, confirming that DeepProfile is capturing key biological processes known to be pivotal in cancer, and paving the way for uncovering more profound insights in subsequent sections of our analysis.

### DeepProfile latent variables capture both cancer and normal tissue-specific expression signatures

We hypothesized that RNA metabolism/ribosomal gene sets were not necessarily identified by DeepProfile because they captured variance related to the presence of different disease subtypes within a tissue of origin, but rather because they contained genes that are constitutively expressed in a highly correlated manner. To test this hypothesis, we generated DeepProfile embeddings for normal tissue gene expression profiles from the GTEx database^27^ (**Supplementary File 1**). By fitting predictor models to differentiate between normal and cancer embeddings, we generated a score for each DeepProfile latent variable denoting how successfully it can separate cancer from normal tissue (**see Methods**). Using DeepProfile pathway-level latent variable attributions, we mapped these latent variable-level scores to pathways to define a cancer-relevance score for each pathway (**see Methods**). A high cancer-relevance score indicates that the pathway is specifically important for cancer because it shows stronger expression variance in cancer than in normal tissues (**Fig. 4 iv and Supplementary File 3**). We found that in comparison with cell cycle pathways, the ribosomal gene sets’ cancer-specificity score was indeed lower (average cancer-specificity score of 82.39 for cell cycle compared to 63.19 for ribosomal pathways; p-value: 1.6 × 10^−17^, Welch’s t-test), indicating that these genes also capture significant variance across normal tissue gene expression profiles. Nonetheless, we note that the degree of biosynthetic activity (as reflected by ribosomal protein expression) has recently been shown to be associated with differentiation state in colorectal cancer^48^, raising the intriguing possibility that DeepProfile’s capture of ribosomal genes reflects variance in differentiation states across tumor samples within a given tumor type. This may explain why some relatively narrowly defined (and therefore more homogeneous) cancer types such as AML did not show significantly enriched ribosome-related pathways. We further note that the two near-universally important pathways with the highest cancer-relevance scores were related to protein folding (prefoldin) and focal adhesions (**Fig. 4**). The latter result is consistent with DeepProfile capturing variation in epithelial-to-mesenchymal transition states that may exist among tumors but would not be expected to occur in healthy tissues.

### Cancer type-specific genes and pathways define molecular disease subtypes

After studying genes and pathways that DeepProfile considered universally relevant, we aimed to identify genes that only capture variance in specific cancer types. We calculated a per-gene cancer type specificity score, defined as the difference between the gene percentile score for one cancer type and the highest gene percentile score across all other cancer types (**Supplementary File 6**).

High specificity scores indicate that a gene captures a large amount of variance in one cancer type but plays a more subordinate role in others (**see Methods**). We found that genes with high specificity scores generally defined dominant *subtypes* or *grades of differentiation* within a tissue category (**Fig. 5a**). For example, the top breast-specific transcripts were prolactin-induced protein (*PIP*), a gene predominantly expressed in well-differentiated estrogen receptor-positive tumors^49^; *FOXC1*, a gene expressed in basal-like breast cancer^50^; and *GFRA1*, which is specific to the luminal A subtype^51^ (**Fig. 5a**).

**Fig. 5.**
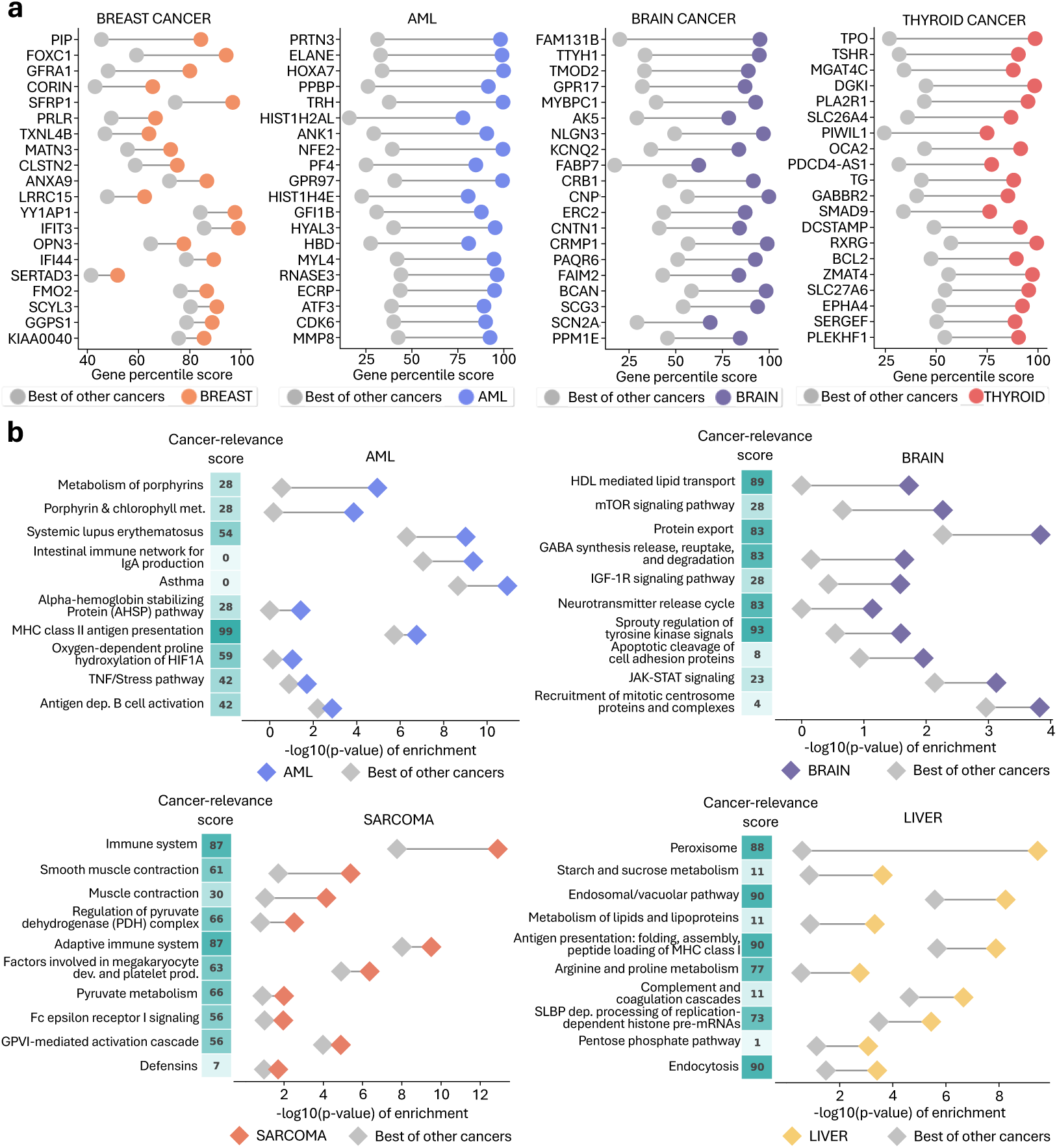
DeepProfile cancer specificity analysis. **a.** The plots of cancer-specific genes shown for 4 cancer types. The difference between the percentile score for the specific cancer type and the highest percentile score among all the other 17 cancer types for the top 20 genes with the highest difference score are shown for each cancer type separately. The colored dots show the percentile score of one gene for the specific cancer type and the gray dots show the highest percentile score the same gene has among all the other cancer types. The genes are sorted based on the difference values. The gene percentile scores for all cancers are available in **Supplementary File 3**. **b,** The plots of cancer-specific pathways along with cancer character scores for 4 cancer types. Pathways are sorted based on the difference between the -log10(p-value) for the specific cancer type and the highest -log10(p-value) among all the other 17 cancer types. Each dot pair represents the -log10(p-value) corresponding to one pathway for the specific cancer type and the highest -log10(p-value) among all the other cancer types. The vector of cancer character scores shows the cancer character percentile score of the latent variable that is capturing the shown pathway. A higher cancer character score indicates that the given latent variable, therefore pathway, specifically important in cancerous tissue. The pathway enrichment scores for all cancers are available in **Supplementary File 3**.

To formally test the hypothesis that DeepProfile captured genes that are differentially expressed among breast cancer subtypes, we calculated the overlap between breast cancer-specific genes and PAM50, a gene set that effectively distinguishes between basal-like, normal-like, luminal A, luminal B, and HER2-enriched subtypes^52^ and obtained significant results (P = 3.8 × 10^−3^) (**see Methods**). Importantly, a linear model (PCA) could not effectively select subtype-specific genes (p-value: 1.0, for PAM50 gene set enrichment), indicating that DeepProfile’s ability to capture non-linear relationships is crucial for learning of biologically meaningful patterns. We further explored the abilities of DeepProfile and traditional linear models (PCA, ICA, RP) to distinguish cancer subtypes, leveraging the Metabric dataset renowned for its detailed subtype labels in breast cancer. The results demonstrated that DeepProfile excels in distinguishing cancer subtypes (**Supplementary Note 3**). However, it is noteworthy that our subsequent analysis also revealed that PCA, despite not efficiently selecting subtype-specific genes, could in fact distinguish between different cancer subtypes. This suggests that while DeepProfile is capable of identifying specific genes tied to cancer subtypes, PCA, with a broader analytical approach, also holds the capability to differentiate between cancer subtypes.

Similarly, AML-specific genes comprised transcripts that had previously been associated with AML subtypes (such as *HOXA7*, *TRH*, *MYL4*, *ANK1*)^53,54^ (**Fig. 5a**) and showed significant overlap with genes identifying AML subtypes (P = 4.2 × 10^−5^)^55^, while PCA again failed (P = 1.0). In the brain, DeepProfile identified genes that distinguish oligodendrogliomas from astrocytomas (such as *CNP*^56^) or vary across glioblastoma subtypes (such as *BCAN*^57^). Top thyroid cancer-specific genes included thyroid peroxidase (*TPO*) and thyroid stimulating hormone receptor (*TSHR*), two transcripts that have critical functions in normal thyroid physiology. These genes may indicate the presence of well-differentiated thyroid cancers, which to some degree retain the expression profiles from their normal tissue of origin, versus highly undifferentiated cancers, which lose tissue-specific transcript expression to a larger degree. To support this hypothesis, we compared DeepProfile thyroid cancer-specific genes with genes associated with thyroid cancer subtypes^58^. We observed that the two gene groups significantly overlapped (p-value: 4.4 × 10^−10^) while the same analysis for the thyroid cancer-specific genes discovered by PCA showed no significance (p-value: 1.0). These case studies demonstrate how DeepProfile successfully detects genes that differentiate cancer subtypes, while a linear model fails to capture these patterns. Cancer-specific genes for each of the 18 human cancers are provided in **Supplementary File 6**.

Next, we extracted curated pathway gene sets that DeepProfile recognized as cancer-specific (**see Methods, Supplementary File 7**). Potentially more informative than a gene-level view, this approach can go beyond categorizing subtype ‘marker genes’ to reveal coherent pathways that vary dominantly among cancers from one tissue of origin. Thus, the analysis provides concrete information about the molecular mechanisms driving expression heterogeneity within cancer types. Indeed, DeepProfile assigned highly characteristic molecular processes to each cancer type.

Top AML-specific pathways were related to porphyrin metabolism and heme biosynthesis (**Fig. 5b**). That leukemic cells show increased heme biosynthesis has been known for more than half a century^59^; but little is known about the mechanistic relevance of the porphyrin production pathway in leukemogenesis. Importantly, recent evidence showing that MYC-overexpressing leukemic progenitors require porphyrin biosynthesis for self-renewal^60^ demonstrates a role for this pathway in driving or facilitating leukemogenesis in a subset of these cancers. It is notable that DeepProfile identified this pathway as relevant to AML, as we are not aware of prior unsupervised analyses that have highlighted porphyrin production. As in our analysis of genes and pathways that were universally important across cancers, we also calculated ‘cancer-relevance’ scores (by comparing matched normal tissue embeddings from GTEx) that determine to what degree a pathway’s importance was specific to malignancy. The AML-specific pathway with the highest cancer-relevance score was MHC class II antigen presentation, represented by *HLA-DMA*, *HLA-DRB1*, *HLA-DMB*, *HLA-DPA1,* and *HLA-DPB1* genes. Downregulated *HLA-DPA1*, *HLA-DPB,* and *HLA-DRB1* in AML has recently been reported during relapse after allogeneic bone marrow transplant and has been interpreted as evidence of graft pressure on leukemic cells^61^. However, DeepProfile’s identification of the MHC class II antigen presentation pathway’s prominence indicates that MHC class II protein expression heterogeneity may be a more general disease feature distinguishing AML subtypes, a concept that has not been described in the literature thus far to our knowledge.

In brain cancer (**Fig. 5b**), lipid transport scored as the most important pathway, with a high cancer-relevance score. Cholesterol is an essential component of myelin, and the brain contains approximately 20% of the body’s total cholesterol^62^. Astrocytes normally produce most of the the brain’s cholesterol, since it cannot be transported across the blood-brain-barrier. In glioblastoma, the brain’s normal lipid metabolism is altered: glioblastoma cells limit cholesterol biosynthesis and depend on exogenous cholesterol uptake for survival^63^, making DeepProfile’s selection of this pathway a notable result. The Sprouty (SPRY) pathway obtained the highest cancer-relevance score, driven mainly by *SPRY1* and *SPRY4*. These two genes negatively regulate FGFR signaling, a pathway that is key to glioblastoma progression and is currently being targeted in clinical trials^64^. These and other examples – such as the identification of an important role for the peroxisome in liver cancer^65^ (**Fig. 5b, Supplementary File 7**) – illustrate DeepProfile’s ability to extract cancer-specific and biologically meaningful expression patterns from large unstructured data depositories. While understanding expression subtypes and the pathways defining them is valuable from a basic science perspective, determining pathways connected to clinical variables is arguably even more important from a translational point of view. We therefore set out to develop a rigorous methodology for connecting DeepProfile embeddings to relevant patient and tumor-level characteristics.

### Detecting survival- and mutation burden-associated pathways via DeepProfile

A pathway’s contribution to DeepProfile latent variables reflects to what degree it captures variance in the primary gene expression data but does not reveal whether the pathway relates to variables of clinical interest. We developed a general methodology for connecting pathways to clinical characteristics via DeepProfile latent variables (**Extended Data Fig. 6** and **Methods**). We tested the approach by extracting pathways that are relevant to two important patient-level and tumor-level features: survival and tumor mutation burden (TMB). Specifically, we associated each DeepProfile latent variable with survival or TMB and generated p-values denoting the association significance between each latent variable and the phenotypes. Then, using the pathway-level attributions for DeepProfile latent variables, we mapped the latent variable-level phenotype associations to pathway-level associations, thereby obtaining survival and TMB association p-values for each pathway (see **Extended Data Fig. 6, Methods**, and **Supplementary Files 8-10**). The same approach can readily be adapted to other variables of interest, for example tumor stage, tumor grade, or treatment response. There are two advantages of using DeepProfile latent variables (instead of genes or pathways themselves). First, as we demonstrated, DeepProfile embeddings encode robust sources of variation among cancer samples; thus, the association search space is reduced to potentially more biologically meaningful variables. These latent variables distill the comprehensive and intricate biological information from the data without relying on predefined features, enabling exploration of relationships with any biological and clinical features. With these latent variables, DeepProfile allows researchers to uncover patterns and associations that might be obscured in the high-dimensional space of gene expression data. Second, since each DeepProfile latent variable is a non-linear combination of genes, it has the unique ability to capture complex interactions between genes and phenotypes of interest. This non-linear mapping allows for the integration of multifaceted biological information, going beyond simple additive effects to model the complex, often non-linear relationships inherent in gene regulation and cellular function. Although these latent variables derived from deep neural networks can offer a more nuanced view, the inherent complexity of these models often complicates interpretation. However, by utilizing XAI methods, we are able to clarify these models, providing interpretable insights that pave the way for the discovery of novel insights into cancer biology.

To test the effectiveness of this approach, we first investigated the curated pathway gene sets that DeepProfile recognized to be significantly related to arguably the most important patient-level trait – survival. As in our previous analyses, we initially focused on pathways associated with survival across all cancer types (**Fig. 6a** and **Supplementary File 11**) (see **Methods**). Notably, in this pan-cancer analysis, the unifying theme of most survival-related pathways was adaptive immunity (**Fig. 6a**). High-scoring gene sets included adaptive immune system, MHC class I antigen presentation, antigen processing cross-presentation, B cell receptor signaling, the proteasome pathway, and activation of NF-**κ**B (all significantly detected in five cancer types). Three pathways stood out for scoring in more than five cancer types. These included DNA mismatch repair (six cancers), a process that can give rise to large numbers of neoantigens when impaired, and MHC class II antigen presentation, which was the highest-scoring pathway overall (significantly detected in seven cancer types). These two pathways will be explored in more detail further below.

**Fig. 6.**
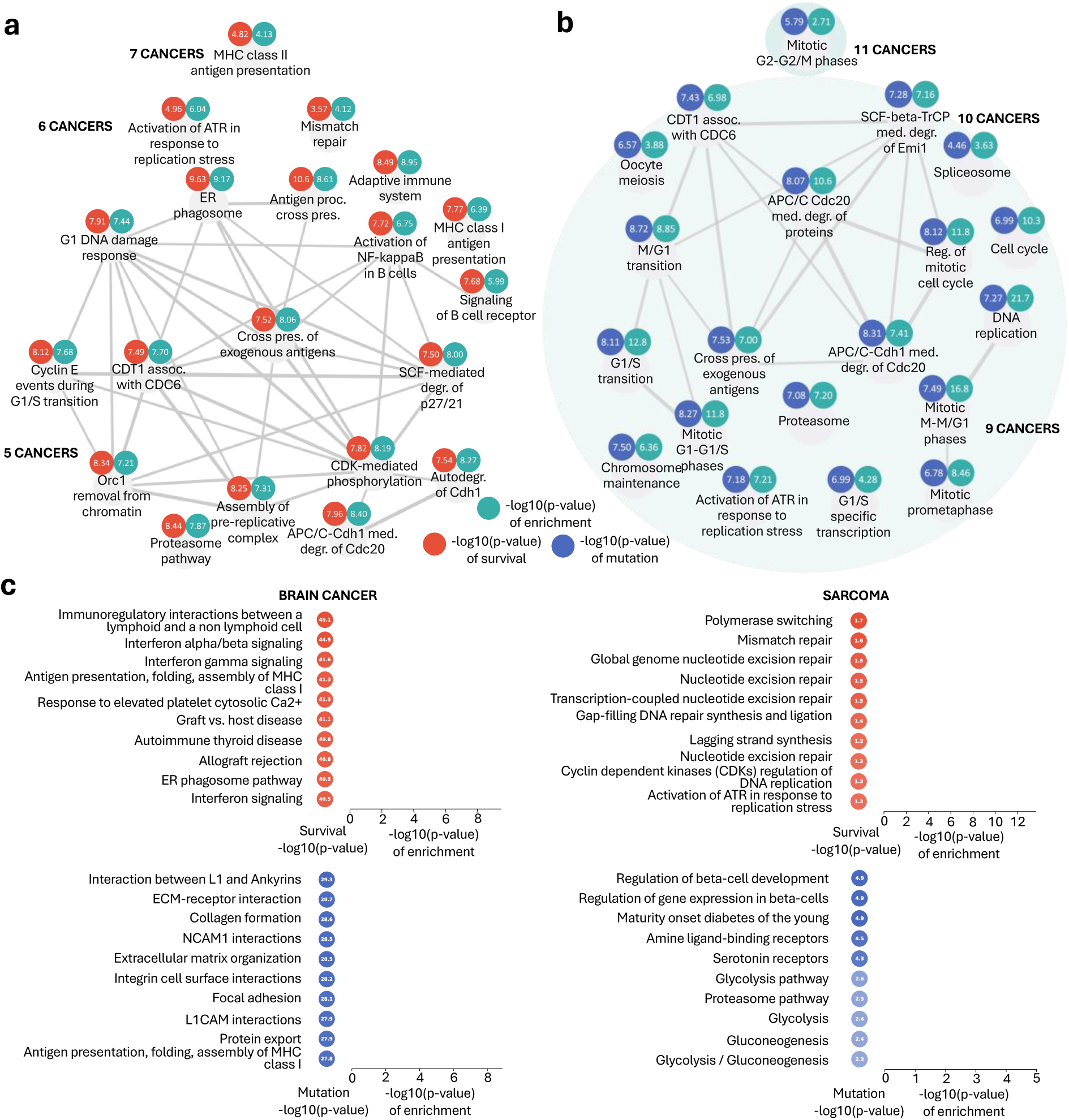
DeepProfile survival and mutation analysis. **(a,b).** The network of top survival-related (**a**) and TMB-related (**b**) pathways. For each pathway group, we show the number of cancers for which the pathway is significantly enriched and significantly associated with survival/TMB (p-value < 0.05). We further show the –log10(p-value) of enrichment and –log10(p-value) of survival/TMB association averaged across all cancers detecting the pathway to be relevant to survival. The connections between pathways are determined based on gene membership Jaccard similarities. **c.** Plots of top survival and mutation associated pathways for brain cancer **(left)** and sarcoma **(right)**. The upper plot shows the top 10 pathways with highest survival scores for the shown cancers along with the survival and enrichment –log10(p-values) and the lower plot shows the top 10 pathways with highest mutation scores for the shown cancers. The scores for all pathways and cancer types are available in **Supplementary File 9**.

To provide a contrast and comparison for these results, we next studied pathways with significant connection to a tumor-level characteristic, TMB (**Fig. 6b** and **Supplementary File 11**) (see **Methods**). Unlike survival, TMB-relevant pathways were most consistently linked to the cell cycle (**Fig. 6b**) and included DNA replication, mitotic M-M/G1 phases, mitotic prometaphase, chromosome maintenance, and others. The top-scoring TMB-linked pathway was mitotic G2-G2/M phases, which was significantly detected in 11 out 18 cancers. These results establish a link between a tumor’s proliferative activity and its mutation burden, consistent with DNA replication acting as a powerful mutagen. This connection carries interesting implications given the strong interest in TMB as a predictor of immunotherapy response^66^.

Analogously to previous analyses, we also studied the pathways with the highest survival and TMB scores for each cancer type. Again, we found that DeepProfile identified distinct sets of pathways as being relevant to both traits. For example, survival-related pathways in brain cancer were dominated by interferon type I and II signaling and MHC class I-mediated immunity, while TMB-related pathways prominently featured cell-cell and cell-matrix interactions (**Fig. 6c**). In sarcoma, survival-related pathways almost exclusively concerned DNA repair processes (mismatch repair, nucleotide excision repair) and replisome function, whereas TMB gene sets were strongly related to glucose metabolism (**Fig. 6c**). Cancer-specific pathway associations with survival and TMB across all 18 cancers can be found in **Supplementary File 8**.

### DNA mismatch repair and antigen presentation via MHC class II are common survival-related pathways

We then explored the unexpected pan-cancer association between survival and DNA mismatch repair and MHC class II antigen presentation in more detail. DeepProfile detects robust correlations between pathways and survival; however, it does not reveal these associations’ directions. Therefore, to define this direction, we fitted univariate Cox regression models on the genes in the pathways being investigated. This returned a survival z-score for each gene and cancer type pair (see **Methods** and **Supplementary File 10**; a negative z-score means that lower expression leads to better chance of survival whereas a positive z-score means that higher expression leads to a better chance of survival).

Examining the z-scores of DNA mismatch repair genes across all cancers, we confirmed a strong correlation with survival (**Fig. 7a**), validating DeepProfile’s findings at the primary gene expression level. The association direction tended to be negative (indicating that lower expression of DNA mismatch repair proteins associates with improved survival), particularly for the six cancers with statistically significant scores in the DeepProfile-based analysis (**Fig. 6a**). We confirmed this finding further using Kaplan-Meier analyses that yielded consistent results (**Fig. 7b** and **Extended Data Fig. 7**) (see **Methods**). The prognostic relevance of DNA mismatch repair gene expression across many cancers is particularly notable given DeepProfile’s identification of the adaptive immune response as a central survival-related pathway hub. Anti-tumor immune responses are thought to depend substantially on the presence of neoantigens, whose abundance increases in cancers with deficient DNA mismatch repair^67^. Similarly, reduced expression of mismatch repair proteins can increase mutability and microsatellite instability^68^. Therefore, higher neoantigen levels in tumors with fewer mismatch repair proteins may make these tumors more visible to the immune system and thus contribute to the improved survival of patients with low DNA mismatch repair protein expression (**Fig. 7c)**.

**Fig. 7.**
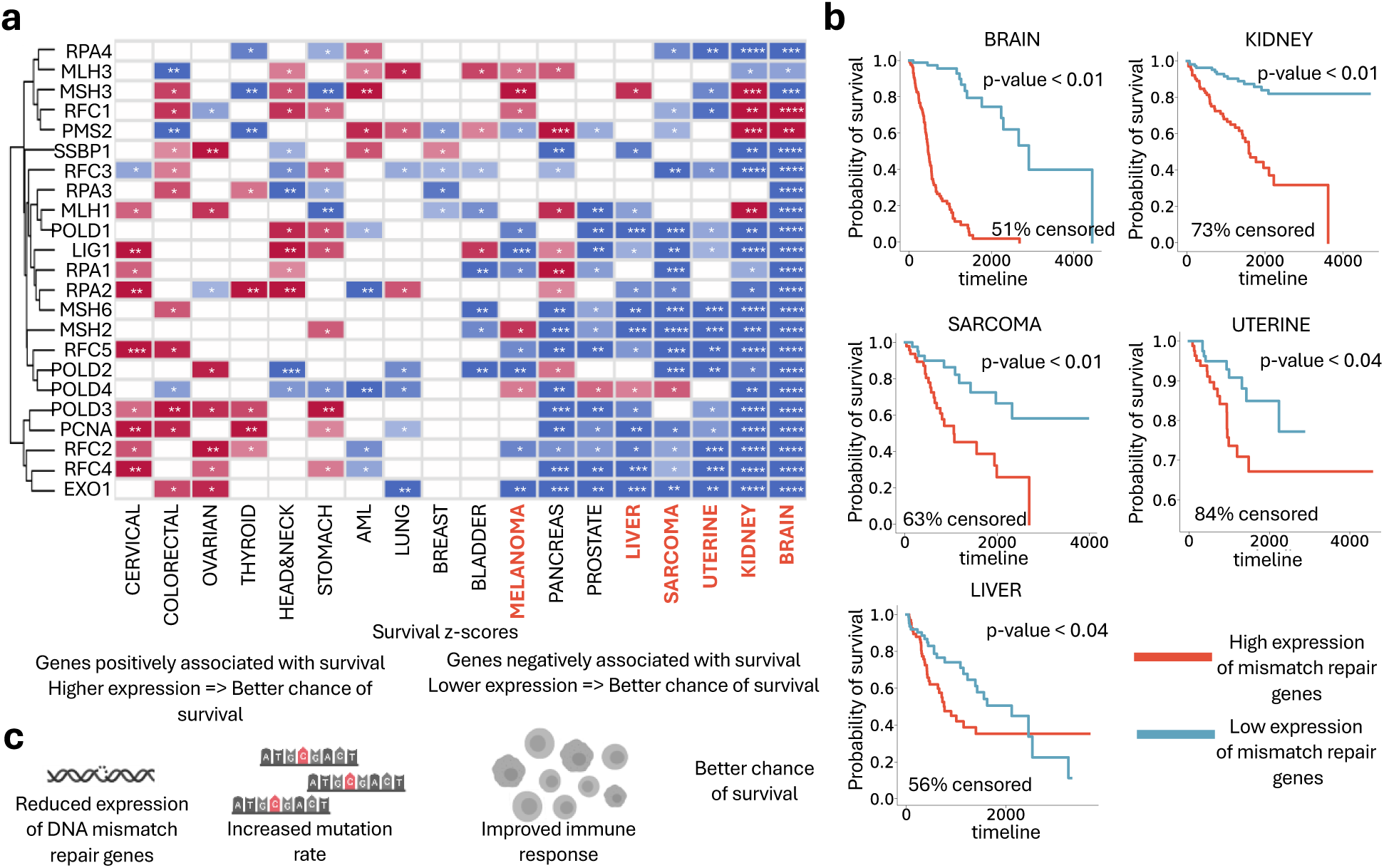
Mismatch repair pathway survival analysis. **a** Heatmap of survival z-scores of all genes included in KEGG mismatch repair pathway (* = magnitude of z-score > 1, ** = magnitude of z-score > 2, *** = magnitude of z-score > 3, **** = magnitude of z-score > 4). 6 cancer types detected by DeepProfile are highlighted. **b.** Kaplan-Meier plots of average expression of mismatch repair pathway. The samples with an expression above (mean + one standard deviation) are marked as highly expressed and below -(mean + one standard deviation) are marked as lowly expressed. The log rank test p-values and the percent of censored samples are reported for each cancer. 5 cancer types with a log rank test p-value below 0.05 are shown. **c.** Schematic of mismatch repair mechanism.

Next, we investigated the MHC class II antigen presentation pathway more thoroughly. We focused on HLA-D genes because they had top-level attribution scores and survival z-scores across all 18 cancer types among all genes in the MHC class II antigen presentation pathway. (The z-scores of all of 91 genes within the MHC class II antigen presentation pathway are provided in **Supplementary File 12**.) Unlike the DNA mismatch repair z-scores, which showed a negative correlation between expression and survival across most cancer types, the association for *HLA-D* expression was bifurcated (**Fig. 8a**). Pancreas, kidney, AML, and brain had a strong negative association between *HLA-D* gene expression and survival change, while the correlation was positive for most other cancers, especially melanoma and uterine cancer. Again, we confirmed these findings via Kaplan-Meier analyses (**Fig. 8b** and **Extended Data Fig. 8**). These results suggested that *HLA-D* gene expression in the tumor and/or its environment is beneficial in some cancer types (melanoma, uterine cancer, breast cancer) and detrimental in others (brain cancer, kidney cancer).

**Fig. 8.**
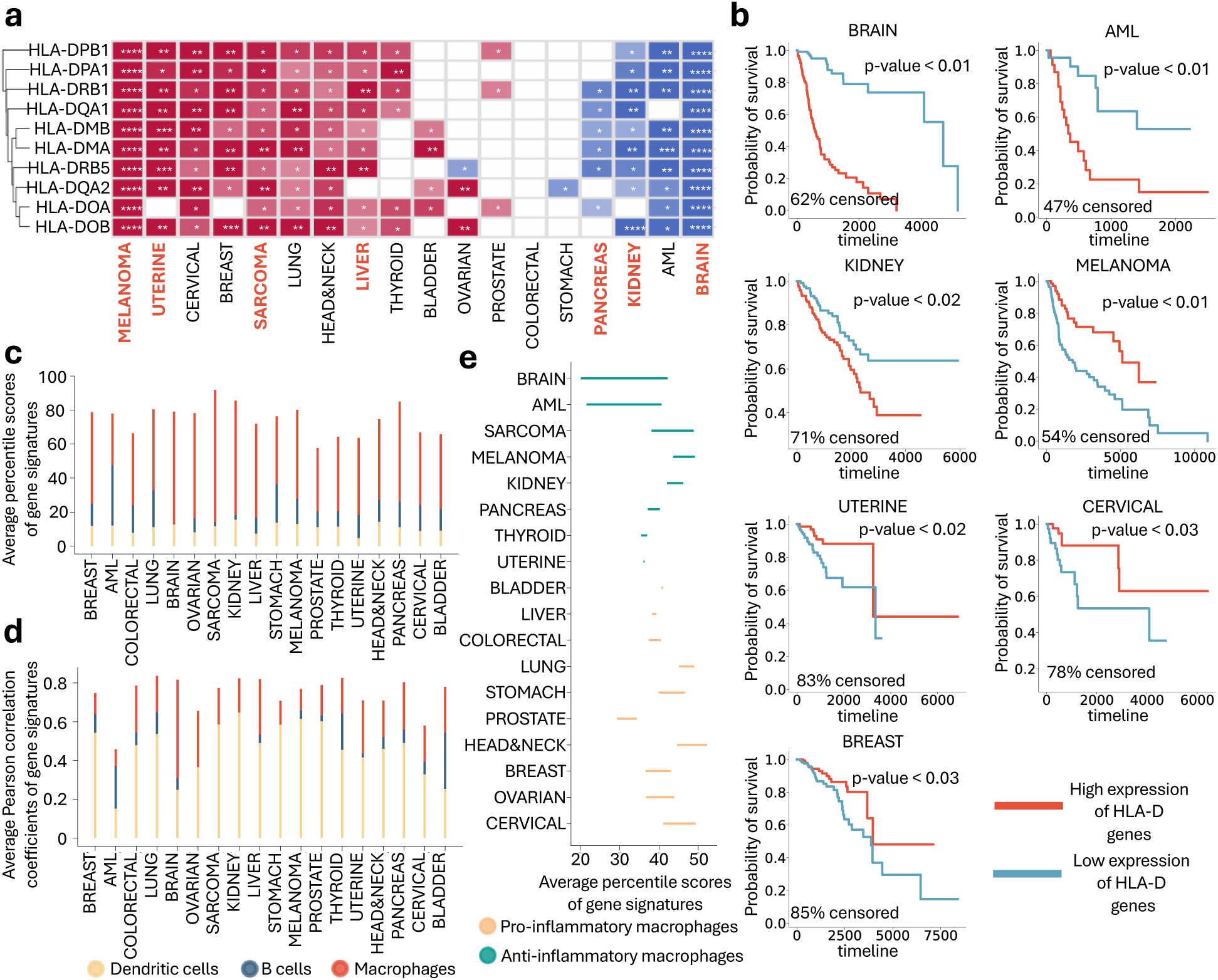
MHC class II pathway survival analysis. **a.** Heatmap of survival z-scores of all HLA-D genes included in Reactome MHC class II antigen presentation pathway. 7 cancer types detected by DeepProfile are highlighted. **b.** Kaplan-Meier plots of average expression of HLA-D genes for cancer types with a log rank test p-value below 0.05. **c.** Comparison of average percentile scores of gene dendritic cells, b cells, and macrophages shown for 18 cancers. **d.** Comparison of average Pearson correlation between the expression of HLA-D genes and cell type signatures for the three cell types shown for 18 cancers. **e.** Comparison of average percentile scores of pro- and anti-inflammatory macrophages shown for 18 cancers. **(See Supplementary Fig. 8)**

Since most cancers do not express MHC class II genes (with the exception of AML, in which *HLA-D* expression is associated with an inflamed phenotype and therapy relapse^61^), we wondered which cell type in the tumor microenvironment might be the primary source of the *HLA-D* transcripts and, by extension, linked to differential survival. Tumor-resident immune cell types that express MHC class II genes include macrophages, dendritic cells, and B cells. To gauge these cells’ relative abundance in the tumor microenvironment, we measured the signature genes’ average percentile score for each cell type, where the most highly expressed gene had a score of 100 (see **Methods**). We found that of the three cell types, macrophage-specific genes were by far the most abundant across all studied cancers, in line with the fact that macrophages can be highly abundant in many cancer types^69–71^ (**Fig. 8c**). Also, we found that in all cancers, the macrophage signature correlated best with HLA-D expression (**Fig. 8d**; see **Methods**), further supporting the notion that macrophages are the largest contributors to *HLA-D* transcript abundance in bulk tumor samples. Considering that macrophages’ divergent functions range from pro-tumor to anti-tumor activity^69–71^, we wondered whether the phenotypes of tumor-associated, *HLA-D-*expressing macrophages might explain the observed bifurcation in the correlation between HLA-D expression and survival. To this end, we examined gene transcripts that may reflect macrophage function. Specifically, we assessed expression of *CD40*, *CXCL9*, *CXCL10*, *CXCL11*, *SLAMF1*, and *TNIP3*, which associate with anti-tumor activity, and of *CFP*, *HRH1*, *NPL*, *PDCD1LG2*, and *CFP*, which typically indicate immunosuppression and tumor promotion^72^. While these genes are not necessarily uniquely expressed by macrophages, the macrophages’ abundance (**Fig. 8c**) makes them plausible main sources of these transcripts. Examining the relative prevalence of the gene transcripts mentioned above revealed that most tumor types expressed both signatures at similar levels (**Fig. 8e**) (see **Methods**). The only large gap, with a large preponderance of immunosuppressive transcripts, was observed in brain cancer and AML – the two cancer types with the most significant negative association between HLA-D expression and survival (P = 3.4 × 10^−2^ and p-value: 1.6 × 10^−1^, Welch’s T test for brain cancer and AML, respectively). We repeated the same test with an extended list of pro- and anti-inflammatory macrophage signatures^73^ and again observed a significantly stronger immunosuppressive macrophage abundance in brain cancer (p-value: 5.0 × 10^−2^ , Welch’s T test) (**Extended Data Fig. 8d**) (see **Methods**). The presence of macrophages that are polarized towards an immunosuppressive phenotype might therefore contribute to the negative correlation between HLA-D expression and survival in brain cancers and AML. In most other cancer types, HLA-D expression correlates with improved outcomes, which is consistent with a net positive effect of macrophages on patient survival.

## Discussion

DeepProfile represents a paradigm for applying unsupervised learning to the analysis of gene expression data. Common unsupervised machine learning techniques in this area fall into three categories: clustering, network inference, and representation learning. The mechanism by which statistical patterns are translated into concrete biological insights is important. DeepProfile represents a major departure from existing unsupervised learning paradigms. While the patterns learned by clustering and network inference algorithms have natural biological interpretations – with gene clusters corresponding to expression modules and network edges corresponding to potential regulatory interactions – representation learning largely lacks methods for such a translation. Linear methods like PCA, ICA, or “shallow” autoencoders have been interpreted by examining the magnitude of their learned weights; however, the “black box” nature of deep neural networks (DNNs) makes it difficult to understand how genes or biological processes are associated with each learned latent variable and how gene expression levels are related to phenotypes. DeepProfile provides a language based on rigorous machine learning principles to “read out” biologically meaningful information from deep representations, enabling discoveries not captured by existing unsupervised analysis paradigms. While DNNs have been successful mainly in tasks where a supervisory label is present^17,74–76^, DeepProfile opens the door for DNN-based approaches to be applied to unsupervised, comprehensive, exploratory analysis of accumulating published gene expression data.

DeepProfile introduces a series of rigorous methodologies to “interrogate” DNNs to generate biological hypotheses. First, one of our key innovations is in the way each latent variable is biologically annotated. We adopted the axiomatic feature attribution method integrated gradients^29^, a principled way of estimating the contribution of each input gene variable onto each latent variable. This enabled the computation of gene importance scores for each latent variable, which can be followed by curated pathway gene sets enrichment analysis on top-scoring genes. Biological characterization of these latent variables is important, for example in cancer, to understand the individual variation in clinical outcomes, response to therapy, and coordinated transcriptional programs underlying cancer progression. The overall gene importance scores computed across all latent variables in the entire model results in top-scoring genes whose expression variation across samples explains a large portion of the expression variation of genes. These genes can be interpreted as master regulators, analogous to “hubs” that are considered important in traditional gene network learning approaches. Additionally, DeepProfile introduces various generalizable methodologies to examine the biological characterization of sample-level phenotypes (such as clinical outcomes and tumor mutational burden) based on the latent variables, the difference between samples with different labels (that is, cancer vs. normal tissues), and differences between different models (that is, cancer types). We showcase DeepProfile’s ability to reveal biological insights through our pan-cancer analysis using these methodologies detailed below.

DeepProfile also introduces a way to ensemble the latent variables from many variational autoencoder models trained using varying numbers of latent dimensionalities and random initializations. The use of integrated gradients^29^ allowed the latent variables of our deep model (**Extended Data Fig. 1**) to be directly ensembled, increasing model stability and consistency, while remaining interpretable. Our experimental results show that DeepProfile’s ensembled latent variables encode general and transferrable information about the cancer transcriptome (**Fig. 2** and **Extended Data Fig. 3**). We also demonstrated that DeepProfile’s ensemble approach can learn better embeddings than individual variational autoencoders trained using specific dimensionalities (**Supplementary Fig. 2**), consistent with the conclusion of Way et al. that models with different latent dimensionality may learn different information^20^. The improvement in performance across a variety of tasks that DeepProfile attains suggests that further studies into ensemble methods for unsupervised gene expression analysis may be fruitful. Furthermore, while DeepProfile was able to extract more underlying biological signal than other unsupervised approaches (**Fig. 2**), the high-dimensional and highly correlated nature of gene expression data means that there may have been more biological signal that was not able to be uncovered. Feature attribution methods tend to split credit among correlated features, potentially “washing out” the signal from large correlated groups^77^. Future work will be necessary to scale methods for disentangling causal effects from observational data to high-dimensional cancer expression data, at the level of either the models or the feature attributions^77,78^.

The application of DeepProfile to a pan-cancer gene expression compendium exposed several intriguing biological patterns. These analyses were enabled by DeepProfile’s integration of the learned model with independent biological databases, including normal tissue expression data, patient level phenotype data, and protein-protein interaction databases. First, we observed that DeepProfile tagged as universally important a very specific category of immune-related genes. Our analysis suggested that these genes did not merely reflect the admixture of different immune cell types in the tumor microenvironment. Instead, they were enriched for cell surface receptors that transduce external signals and thus influence downstream gene expression in a variety of immune cells. Why do these genes capture variance so efficiently? The simplest explanation is that they are representative of recurring transcriptional phenotypes of common immune cells. Depending on the level of immune cell admixture - and thus the magnitude of the immune cell contribution to the overall expression profile - this may be sufficient to propel these genes to such a prominent position. However, an even more powerful explanation is that transcriptional states of malignant cells and infiltrating immune cells are correlated to some degree. For example, cancers with high expression of genes indicative of epithelial-to-mesenchymal transition exhibit a distinct, suppressed immune landscape^79^. Single cell sequencing studies have shown that transcriptional profiles of immune and cancer cells can co-vary and suggest the existence of recurring “hubs” of interacting cells^80^. Genes that are characteristic of such hubs would be expected to capture particularly high levels of variance, as they would be predictive of both immune and tumor cell transcriptomes. Identification of such genes may be of particular interest from a therapeutic perspective. Careful investigation of top universal DeepProfile genes in single cell gene expression data across different cancers will undoubtedly shed more light on this question in the future.

In our cancer-specificity analysis, DeepProfile excelled at extracting disease subtype-specific signatures from the data in an unsupervised manner. We consider this impressive, given that the input datasets were not curated and carefully standardized, such as the ones that were used for the initial discovery of these signatures, but unstructured and variable data deposited in a public database by hundreds of different research groups. DeepProfile’s excellent performance in this setting shows that it can robustly identify relevant biological signals in challenging situations in which other methods (like PCA) do not perform adequately. Analysis of cancer-specific DeepProfile pathways identified disease-specific processes, such as porphyrin metabolism in AML or lipid transport in brain cancer. By further annotating these pathways by their specificity to malignancy, highlighting those that play a comparatively minor role in normal tissue gene expression (via embeddings of GTEx profiles), DeepProfile has generated a list of prime candidate pathways that can be explored for therapeutic intervention opportunities.

Perhaps the most interesting aspect of our analysis was the establishment of a quantitatively rigorous connection between DeepProfile embeddings and patient survival characteristics. The results were unexpected and surprising. Low expression of DNA mismatch repair transcripts was significantly associated with improved survival in this large cohort of varied cancer types, most of which are expected to be mismatch repair proficient. These results suggest that capacity for DNA mismatch repair may exist on a transcriptionally-driven spectrum and that a tumor’s exact position on this continuum may be therapeutically relevant. Microsatellite unstable tumors across all tissues respond well to immune checkpoint therapy, and are thus universally approved for treatment with pembrolizumab^81^. Our results raise the question whether cancers with low DNA mismatch repair gene expression might also benefit from immune checkpoint inhibition.

Finally, analysis based on DeepProfile’s latent spaces showed that adaptive immunity pathways, particularly those related to MHC class II antigen presentation, were the most consistently survival-related among 1,077 tested functional gene sets, the latter surpassing even DNA mismatch repair. This surprising result was highly specific to patient survival, as demonstrated by a comparative analysis for TMB, in which the adaptive immune system did not play a significant role. Focusing on the top-scoring genes from the MHC class II antigen presentation gene set, we found that HLA-D transcripts were largely responsible for the strong outcome association. Given that a limited number of immune cells express HLA-D genes, we were able to nominate macrophages as the ‘prime suspect’ source of these survival-associated transcripts in the tumor microenvironment. The effect of HLA-D expression, however, was bifurcated across tumor types. Brain cancer and AML patients had a worse outcome if HLA-D expression was high, while melanoma and uterine cancer patients benefitted. We speculate that the transcriptional phenotype of tumor-resident macrophages (pro- or anti-inflammatory) determines whether the presence of these cells has a net beneficial or harmful effect. We found that in glioblastoma, expression of transcripts characteristic of anti-inflammatory macrophages, which are thought to drive tumor progression^82^, was predominant, potentially explaining the negative correlation between HLA-D expression and outcome. Pro- and anti-inflammatory macrophage transcripts were more balanced in other tumor types, including melanoma and uterine cancer. In these cases, the net effect of the total macrophage population appears to be positive. Importantly, these results are in line with a recent meta-analysis which suggested that expression of anti-inflammatory macrophage markers was correlated with worse prognosis across multiple cancer types, while expression of pro-inflammatory markers was associated with improved survival^82^. Again, it will be important to follow up on these observations in single cell data sets, once their size has grown sufficiently to conduct robust survival analyses, or in more extensive immunohistochemical studies of macrophage polarization across large patient cohorts.

In summary, we have devised and implemented a deep learning framework to extract robust biological signals from large-scale cancer gene expression data. DeepProfile is designed to be a resource for the cancer research community. Using our framework, researchers can create robust and interpretable embeddings of new expression data (**Extended Data Fig. 2**), improving performance on downstream tasks and increasing insight into relevant transcriptional programs in their samples. The demonstrated compatibility between microarray data and bulk RNA-seq data (**Extended Data Fig. 3**) suggests that the learned model can be used for bulk RNA-seq data as well. Beyond the computational advance represented by this approach, DeepProfile provides hundreds of biological insights gleaned from existing compendia that can be mined by researchers to advance our understanding of different human malignancies.

## Acknowledgements

We thank Sara Mostafavi and Sheng Wang for careful review of the manuscript, Nicasia Beebe-Wang and Nao Hiranuma for helpful comments on our experiments, and Kaley Joyes for help with editing the manuscript. This work was funded by the National Science Foundation [DBI-1759487, DBI-1552309 to SIL]; American Cancer Society [127332-RSG-15-097-01-TBG to SIL]; the Mark Foundation for Cancer Research (to KN), the American Association for Cancer Research (to KN) and the National Institutes of Health [R35-GM128638 to SIL and R37-CA225655 to KN]

## Notes

### Competing Interest Statement

The authors have declared no competing interest.

### Summary of Updates

All figures revised. Supplemental files updated.

