## Supplementary Methods for "A deep profile of gene expression across 18 human cancers"

### KEY RESOURCES TABLE

| REAGENT or RESOURCE | SOURCE | IDENTIFIER |
|---|---|---|
| **Software and algorithms** | | |
| Tensorflow Keras | https://github.com/keras-team/keras | |
| GEOParse | https://github.com/guma44/GEOparse | |
| ComBat Python | https://github.com/brentp/combat.py | |
| Scikit-learn | https://github.com/scikit-learn/scikit-learn | |
| GMeans | https://github.com/flylo/g-means | |
| Integrated Gradients | https://github.com/hiranumn/IntegratedGradients | |
| SciPy | https://github.com/scipy/scipy | |
| Cytoscape | https://cytoscape.org/ | |
| survival : Survival Analysis | https://cran.r-project.org/web/packages/survival/index.html | |
| lifelines | https://github.com/CamDavidsonPilon/lifelines | |

### RESOURCE AVAILABILITY

#### Lead contact

#### Materials availability

#### Data and code availability

The input gene expression datasets, their lower-dimensional embeddings, gene-level and pathway-level relevance, and the results of our pan-cancer analysis are publicly available at: https://github.com/suinleelab/deepprofile-study (code), and https://doi.org/10.6084/m9.figshare.25414765.v2 (data).

### EXPERIMENTAL MODEL AND SUBJECT DETAILS

**Not applicable.**

### METHOD DETAILS

We downloaded publicly available gene expression datasets generated by either of the two microarray platforms: Affymetrix GeneChip Human Genome U133 Plus 2.0 (Affy HG-U133 Plus 2.0) and Affymetrix GeneChip Human Genome U133A 2.0 (Affy HG-U133A 2.0). These datasets were available from the National Center for Biotechnology Information (NCBI) Gene Expression Omnibus (GEO) database[1] for 18 cancer types and we used the *GEOparse* Python library (https://github.com/guma44/GEOparse) for downloading the datasets. A list of GEO search keywords, downloaded series, and number samples and genes for each cancer type is available in **Supplementary File 1**.

While GEO searching filters results according to supplied keywords, the returned results may still include gene expression samples from healthy tissues or patients with cancer types other than the queried cancer type. To eliminate these irrelevant samples, we removed the samples that do not contain the search keywords in their *titles*, *characteristics*, or *descriptions*. To further clean our data without unnecessarily eliminating relevant samples, we manually curated it. Using these steps, we aimed to minimize the number of incorrectly included and incorrectly excluded samples. We also excluded cell line expression samples and used only patient samples because the same cell line's low expression variance across datasets might prevent deep neural networks from learning a reliable model. Despite our automated and manual curation to eliminate samples from cell lines, other cancer types, and healthy tissue, it is still possible that some outlier samples are included in our GEO data collection.

To integrate data from various platforms, we converted platform-specific probe IDs to gene symbols using the probe ID to gene symbol conversion lists for each platform available in GEO.

For each cancer, we took the genes present in all data series we have available. A study might have different sample batches submitted on different dates indicated in the *submission_date* field. We corrected for these potential batch effects within each study using the Python ComBat[2] library's *combat* function with the default parameters (https://github.com/brentp/combat.py), where different batches correspond to data subsets submitted at different dates. We log transformed the expression measurements, standardized (i.e., zero-mean and unit variance) each gene in each dataset to ensure that different input features (i.e., gene expression levels) are on the same scale, and applied mean imputation to impute missing gene-level measurements. We also excluded duplicate samples with the same GEO IDs. We concatenated all datasets and applied batch effect correction, once again using ComBat with the same parameters, considering each study to be a separate batch in order to minimize the effect of potential study-specific confounders.

**Training variational autoencoder models**

An *autoencoder* is a type of neural network that consists of an encoder and a decoder network with an information bottleneck layer with $D$ latent variables (i.e., $D \ll M$) in the middle[3]. It generates an embedding $Z$ such that the information present in the original space is preserved in this lower dimensional space as well. Specifically, the encoder network, defined as $f_{\emptyset} : X \longrightarrow Z$, maps from the input space $X \in \mathbb{R}^M$ to latent embedding $Z \in \mathbb{R}^D$. Similarly, the decoder network, defined as $g_{\varphi} : Z \longrightarrow X$, maps the embedding $Z$ back to input space. We optimize over the both networks to minimize the squared 2-norm distance between our input $X$ and the reconstructed input as follows:

$$min_{\phi,\varphi} \mathbb{E} \parallel x - g_{\varphi}( f_{\emptyset}(x)) \parallel_2^2.$$

A *variational autoencoder* (VAE) is an extension of a standard autoencoder that takes as input an $N \times M$ matrix $X$, where $N$ denotes the number of samples, $M$ denotes the number of features and $X_{ij}$ denotes the feature $j$ of sample $i$. It also consists of encoder and decoder networks but adopts a regularized training such that the model is robust to overfitting[4]. To perform regularization, VAE learns a distribution of the latent space rather than learning the encoding directly and samples from the learned distribution to generate an embedding. VAE trains the model to bring the distribution of the latent space as close to a standard Gaussian distribution (i.e., N (0, 1)) as possible, which ensures that the learned distribution is regularized.

We define the encoder network as $f_{\emptyset} : X \longrightarrow \mu_x, \sigma_x$, which maps from the input space $X \in \mathbb{R}^M$ to latent space distribution mean $\mu_x \in \mathbb{R}^D$ and distribution variance $\sigma_x \in \mathbb{R}^D$. We then sample from the distribution to define the low-dimensional embedding $Z \in \mathbb{R}^D$:

$$Z \sim N(\mu_x, \sigma_x).$$

The decoder is defined the same way as it is in a standard autoencoder. To regularize the distribution over the latent space, VAE adds a regularization term to the model's loss function, i.e., Kullback-Leibler divergence between the learned distribution and a normal distribution[5]. The network is trained to be optimized as follows:

$$min_{\phi,\varphi} \mathbb{E} \parallel x - g_{\psi}( f_{\emptyset}(x)) \parallel_2^2 + KL[(\mu_x, \sigma_x), \mathrm{N}(0, 1)],$$

where $KL[(\mu_x, \sigma_x), \mathrm{N}(0, 1)]$ denotes the Kullback-Leibler divergence between the distributions. This regularization component forces the encoder and decoder networks to learn a generalizable, smooth latent space that embeds similar samples close to each other.

Before training different VAE models for a cancer type, we extracted the principal components[6] of the expression matrix; we trained the VAEs using these components as inputs, a commonly used approach for training deep neural networks to prevent overfitting[7]. We chose the number of principal components based on the number of samples and their ability to explain a significant portion of the variance in the data (see **Supplementary File 1** for the number of components for each cancer type). Specifically, we selected 1,000 components for cancer types with more than 1,000 samples, 500 components for those with 500 to 1,000 samples, and 250 components for those with fewer than 500 samples. Our criteria ensure that the selected components account for approximately 80% of the variance in almost all cancer types and 90% for most (**Supplementary Table 2**).

We trained VAE models using the principal components of the cancer-specific gene expression matrix as inputs; the encoder and decoder networks both include 3 fully connected layers, and the two networks mirror each other in structure. The minibatch size is set to 50, and we trained the models using the Adam optimizer[8] with a learning rate of 0.0005. We initialized each VAE model with a different random set of weights using *Glorot_uniform* weight initialization. We built the entire model in Python using *Keras* with *Tensorflow* backend (https://github.com/keras-team/keras).

In determining the size of the latent space for our VAE models, we specifically selected a set of sizes - 5, 10, 25, 50, 75, and 100. This deliberate selection was made to give our models a broad scope to capture a comprehensive range of information from the data. We established these sizes to provide a structured approach to encompass the variety and complexity of the data patterns we are analyzing. All layers use rectified linear unit activation except the last layers of both networks, where we applied linear activation and batch normalization on all encoder layers. Additionally, we fine-tuned the VAE models' hyperparameters, including the dropout rate and the number of neurons per layer, utilizing 5-fold cross-validation and gauging the fine-tuning by the metric of validation reconstruction error. Our options for dropout rate included 0, 0.2, 0.4, and 0.6. Regarding the number of latent variables in the intermediate layers, we considered configurations such as (50, 5), (100, 25), (250, 50), (250, 100), and (300, 150).

Initially, we calibrated the dropout rate by averaging the validation reconstruction errors across all models with different configurations of intermediate layers, particularly highlighting the findings for breast cancer (which has the largest sample size), sarcoma (with a sample size that represents the average), and bladder cancer (with the smallest sample size) in **Supplementary Fig. 4A-C.** Models set with a dropout rate of 0 showed the lowest average reconstruction errors. Fixing the dropout rate at zero, we then optimized the count of latent variables in the intermediate layers. The results, depicted in **Supplementary Fig. 4D-F**, show that models with an increased number of neurons exhibit improved performance. Nevertheless, to ensure a balance between the efficiency of model training and the precision of feature importance assessments, we settled on 250 and 100 latent variables for the first two layers for latent space sizes of 25, 50, 75, and 100. For a latent space size of 10, the numbers were 250 and 50 latent variables, and for the size of 5, 100 and 25 latent variables were selected. We show the training and validation loss across different latent dimensions in **Supplementary Fig. 5**.

The GPU memory usage for the VAE models of DeepProfile is detailed in Supplementary Table 3. The maximum GPU memory usage documented across 18 different cancers is 475MB, demonstrating the efficiency of DeepProfile framework in handling large-scale genomic data.

**Learning DeepProfile latent variables**

DeepProfile combines all embeddings generated by VAE models to learn a single, robust latent space that can preserve both high- and low-level features. We trained a total of |D|*|R| models, where D is a set of possible latent space sizes for individual VAE models and R is a set of random seeds used to initialize model weights. We trained a VAE model for each latent space size $d \in D$ and for each random seed $r \in R$ for the initial weights, which we denote as $VAE_{d,r}$. For our experiments, we used D = {5, 10, 25, 50, 75, 100} and R = {0, …, 99}), which corresponds to 100 random models for each of the 6 latent space sizes, for a total of 600 VAE models. Each VAE model takes the expression matrix $X \in \mathbb{R}^M$ as input and outputs an embedding $Z \in \mathbb{R}^d$. We assessed the robustness of the DeepProfile model by analyzing gene ranking consistency across multiple VAE model ensembles, as detailed in **Supplementary Note 4**. This analysis showed a substantial increase in gene ranking overlap as more models were added to each ensemble, demonstrating the model's robustness and stability.

Across all |D|*|R| models, we have |D|*|R| embeddings and $\sum_{d \in D} \mathrm{d} * |\mathrm{R}|$ latent variables in total (600 embeddings and 26,500 latent variables for our setting). To group similar data encodings, we applied k-means clustering to cluster all latent variables from all models. We used the Python *sklearn* library's *KMeans* model with k-means++ initialization and 10 different starting points[9]. k-means assigns each of $\sum_{d \in D} \mathrm{d} * |\mathrm{R}|$ latent variables to one of the L clusters, where L is the number of DeepProfile latent variables. Note that we disregard the information about which latent variable came from which model: we simply applied clustering to all latent variables by treating them as independent and identically distributed (i.i.d.). As a result, different latent variables of the same VAE model might be in different clusters as well as in the same cluster. Also, one cluster may include latent variables from different models with the same latent space size (i.e., different runs), or it can also include latent variables from models with different latent space size. After k-means groups the latent variables that are similar across runs and dimensions, we created one ensemble latent variable per cluster by averaging the values of all latent variables in that cluster to obtain a final embedding, $Z \in \mathbb{R}^L$ (**Extended Data Fig. 1 and 2a**).

To select the latent embedding size for DeepProfile, we applied *G-means clustering*, an extension of k-means clustering that determines the optimal number of clusters k[10]. We used Python's *gmeans* package and trained with strictness criteria 3, maximum depth 10, and minimum observation count 1 (https://github.com/flylo/g-means). For each cancer type, we fitted G-means clustering before training the k-means models to select the optimal k value. We averaged the optimal number of clusters across 18 cancers to set L = 150 as the final latent embedding size after rounding down the exact average, which was 157. We selected the same latent size for each cancer type to enable direct comparison between cancer-specific embeddings. To address the inherent variability in k-means clustering, particularly in initial centroid selection, we performed stability analyses, detailed in the **Supplementary Note 5**. These analyses, involving Normalized Mutual Information (NMI) scores and gene/pathway comparisons across runs, consistently affirmed the stability of our model in identifying key genetic elements.

Our DeepProfile framework can encode user cancer expression samples. When user expression samples are passed to the DeepProfile model, we first apply the same preprocessing procedure we applied to our training samples after eliminating the genes not available for the training samples. We pass the preprocessed expression matrices to our trained VAE models to generate embeddings. In other words, we use the learned weights for VAE models to encode the user samples and generate an embedding from each VAE model. We then use the learned ensemble assignments to cluster VAE latent variables and take the average value in each cluster to define the final DeepProfile embedding for user samples (**Extended Data Fig. 2b**). Users can select the number of latent dimensions, in which case ensemble label assignments will be calculated again to define the new ensemble latent variables for the user-selected latent dimension size.

We provide the estimated training and testing times for all cancer types in **Supplementary Table 4**. On average, the average training time for each cancer type is approximately 1.36 hours, and the average testing time is notably efficient at just 0.10 hours. This indicates that while the training phase of the DeepProfile models requires a reasonable amount of time, the testing phase is exceptionally efficient, which is advantageous for practical applications.

### Gene- and pathway-level attributions of DeepProfile latent variables

To calculate gene-level attributions of DeepProfile latent variables, which denote how much each gene contributes to the learned latent variables, we used Python's *Keras* implementation of Integrated Gradients (https://github.com/hiranumn/IntegratedGradients), a gradient-based feature attribution method for neural networks[11] **(Extended Data Fig. 2c)**. When applied to a neural network model, Integrated Gradients learns the sample-level importance values of each input feature for each output variable.

In order to compute the gene importance values for each latent variable in our finalized DeepProfile model, we follow a two-step approach. Firstly, we calculate the Integrated Gradient (IG) values for each principal component relating to every Variational Autoencoder (VAE) latent variable. Subsequently, these IG values are multiplied by the respective principal component weights, also known as eigenvectors. The process of multiplying the IG values by the eigenvectors provides a mechanism for scaling the importance values according to the influence of each principal component on the original genes. Thus, we are able to obtain the gene importance values linked to each VAE latent variable. As the DeepProfile model is an ensemble of VAE models, the DeepProfile latent variables include numerous VAE latent variables. Therefore, importance values for each DeepProfile latent variable are calculated by averaging the attributions of the corresponding VAE latent variables.

To determine the global importance of each gene for a latent variable, we calculate the absolute valued average of attribution scores across all training samples for each cancer type. Since DeepProfile is an ensemble of VAE models, where each DeepProfile latent variable combines multiple VAE latent variables, feature attributions for each DeepProfile latent variable are calculated by averaging the attributions of the VAE latent variables defining that ensemble latent variable.

To calculate pathway-level attributions, we used gene-level attributions and ran pathway enrichment tests using a total of 1,077 functional pathways from Reactome[12], BioCarta[13], and KEGG[14] from the C2 collection of the version 6.2 of MSigDB[15,16] . For enrichment tests, we used Fisher's Exact Test's (FET)[17] *fisher_exact* method from Python's *scipy.stats* module. From the gene list for each pathway, we removed the genes that are not present in our input expression matrix and passed the top G genes with the highest importance values for a DeepProfile latent variable to FET, where G is the average pathway length across all 1,077 functional pathways from Reactome, BioCarta, and KEGG. For multiple hypothesis correction, we applied Benjamini-Hochberg FDR correction[18] across all latent variables, using the *multipletests* function from Python's *statsmodels* library.

**Comparing DeepProfile to alternative dimensionality reduction methods**

We compared DeepProfile to alternative dimensionality reduction algorithms, including the commonly used linear methods as well as other deep learning approaches. We trained these algorithms using the same preprocessed gene expression levels that we used as input to DeepProfile VAE models.

**Gaussian random projection** maps the original input to a lower dimensional space, where each component is randomly drawn from a normal distribution. From Python's *sklearn* library, we used *GaussianRandomProjection* and repeated training 10 times with different random seeds to output 10 different embeddings.

**Principal Component Analysis (PCA)**[6] is a linear dimensionality reduction method that generates orthogonal components to encode variation in the original input space. We used the *PCA* module from Python's *sklearn* library and used the top 150 principal components when comparing it to the DeepProfile embedding, which has 150 latent variables.

**Independent Component Analysis (ICA)**[19] is also a linear dimensionality reduction method that learns independent components from the original space. We trained ICA using Python's *sklearn*

*FastICA* with 100,000 iterations; we repeated the training 10 times with different random seeds to output 10 different embeddings.

**Autoencoder (AE)**[3] is a deep unsupervised neural network consisting of an encoder and decoder network trained to learn a latent space that can reconstruct the original space as successfully as possible. For autoencoder trainings, we used the same top principal components of the preprocessed gene expression levels as we did for training DeepProfile to enable a fair comparison between models. We tuned the hyperparameters of AE models, the number of layers, number of latent variables, dropout rate, and batch size using 5-fold cross validation with reconstruction error as the metric. In the final AE model, we have 1 hidden layer each in encoder and decoder networks with 750 latent variables, 0.1 dropout rate, and batch size of 100. The model was trained with the Adam optimizer using a learning rate of 0.0005. Since each different random initialization of the model can output a different representation, we repeated the autoencoder training 10 times with different random weight initializations. The models were implemented using the *Keras* with *Tensorflow* backend.

**Denoising Autoencoder (DAE)**[20] is a regularized autoencoder model that adds noise to the input data in order to generate more robust embeddings. We applied the same procedure to denoising autoencoder models as autoencoders: we passed the same preprocessed gene expression levels as input to DAE models and selected the hyperparameters with 5-fold cross validation. The final tuned model has 1 hidden layer each in encoder and decoder networks, 750 latent variables and 0.1 dropout rate. We optimized the model using the Adam optimizer with a learning rate of 0.0005 and batch size of 100. We again repeated the training of DAE models 10 times with different random weight initializations. The models were implemented using the *Keras* with *Tensorflow* backend.

**Variational Autoencoder (VAE)** We included the single VAE models with 100 latent variables — the most powerful configuration among all models within our DeepProfile ensemble — as a baseline for comparison. The VAE models have two hidden layers, featuring 250 and 100 latent variables respectively. The dropout rate is set to 0. Optimization is achieved using the Adam optimizer at a learning rate of 0.0005 and a minibatch size of 50.

**Creating TCGA RNA-Seq embeddings**

We downloaded TCGA RSEM normalized log2 transformed RNA-Seq expression matrices for all cancer types from Broad Institute data version 2016_01_28 (https://gdac.broadinstitute.org/) and generated by TCGA Research Network (https://www.cancer.gov/tcga/). The mapping of DeepProfile and TCGA cancer types as well as the number of samples are listed in **Supplementary File 1**. We preprocessed the TCGA expressions with the same pipeline we used for preprocessing GEO expression datasets: we selected the genes available only in the training data, zero imputed the genes missing in the TCGA dataset, and standardized each gene to zero-mean univariance.

Since we take the top principal components of the training data to train DeepProfile, we applied the same processing step for generating TCGA embeddings. We encoded the TCGA samples using the PCA model trained on the training data. To generate the DeepProfile embeddings, we loaded all trained VAE models, encoded TCGA PCA transformed input features with each of the models, and used the pre-learned ensemble labels to cluster the latent variables of our VAE embeddings and define a 150-dimensional DeepProfile embedding for TCGA RNA-Seq samples. We repeated this procedure for each cancer type. To assess the generalizability of the DeepProfile model to TCGA data, we measured the Mean Square Error (MSE) on both GEO and TCGA datasets across different latent dimensions. Significantly lower MSE values on TCGA data, as detailed in **Supplementary Fig. 6**, underscore DeepProfile's proficiency in effectively reconstructing and adapting to unseen data, demonstrating its robust generalization capabilities.

Similarly, for all the alterative dimensionality reduction approaches, we used the trained models to encode TCGA RNA-Seq samples.

**Comparison of DeepProfile microarray and RNA-Seq embeddings**

To demonstrate that DeepProfile can learn informative latent spaces from both microarray and RNA-Seq test data, we used the TCGA cancer samples for which we have both RNA-Seq and microarray expression available. We downloaded TCGA log2 LOWESS normalized microarray expression matrices for all the available cancer types from Broad Institute data version 2016_01_28 (https://gdac.broadinstitute.org/) generated by the TCGA Research Network (https://www.cancer.gov/tcga/) (see **Supplementary File 1** for the mapping of cancer types and the number of samples for which we have matching expression measurements). We selected the genes present in both microarray and RNA-Seq datasets to enable a fair comparison and preprocessed microarray expression profiles following the same preprocessing steps applied to GEO samples. We then measured the Pearson correlation coefficient between the gene expression matrices generated with the two technologies using the Python *scipy.stats* library's *pearsonr* method. Thus, we obtained a correlation coefficient for each TCGA sample, which denotes the similarity between the two expression profiles.

Following the same procedure for creating DeepProfile TCGA RNA-Seq embeddings, we created DeepProfile embeddings from TCGA microarray profiles. In this way, using the DeepProfile framework, we obtained two separate embeddings for a cancer type: (1) embeddings generated from microarray expression, and (2) embeddings generated from RNA-Seq expression. We then measured the Pearson correlation between DeepProfile RNA-Seq and microarray embeddings for each TCGA sample using the Python *scipy.stats* library's *pearsonr* method. Again, we obtained a correlation coefficient for each TCGA sample, which denotes the similarity between two expression embeddings.

**Comparing DeepProfile pathway coverage to alternative dimensionality reduction methods**

When comparing DeepProfile to other dimension reduction methods in terms of pathway coverage, which we used as a metric for evaluating the biological relevance of the learned latent space, we followed the same procedure as we used for DeepProfile. We applied Fisher's Exact Test (FET)[17] *fisher_exact* method from Python's *scipy.stats* module and obtained a p-value for each latent variable-pathway pair, denoting the significance of enrichment.

To run pathway enrichment tests, we first obtained the gene-level attributions for each dimensionality reduction method. For PCA, we obtained the component matrix, which denotes the contribution of each gene to each principal component, and we took the absolute values of the component matrix for use in enrichment tests. Similarly, for ICA and RP, we obtained the absolute valued component matrices. Since we trained each model 10 times with different random initializations, we repeated the FET for each of the 10 models and averaged the pathway enrichment results over 10 runs. For autoencoder and denoising autoencoder models, we used Integrated Gradients [11] to obtain gene-level attributions for the embedding latent variables, following the same procedure we applied for VAE models. Again, we obtained gene-level attributions for each of the 10 random trainings, conducted FET enrichment tests for each run, and reported average pathway enrichments over 10 models.

We compared DeepProfile's pathway coverages to other dimension reduction methods using 3 different metrics:

1. We compared the *average pathway coverages*. The enrichment tests we conducted provided us with an enrichment p-value for each latent variable-pathway pair. After FDR correction, we marked the latent variable-pathway pairs with a p-value < 0.05 as significant and calculated the total number of significant enrichments for each latent variable. We defined the number of pathways significantly captured by each latent variable as the pathway coverage of that latent variable. Then, we averaged these latent variable-level pathway coverages across all latent variables to calculate the average final coverage of an embedding. This metric let us define an average pathway coverage score per model and per cancer type.

2. We compared the *distributions of latent variable-level pathway coverages* across models. Again, using the same pathway-level attribution p-values, we counted the number of pathways significantly captured (FDR p-value < 0.05) by each latent variable of each embedding. We compared distributions for each method and each cancer type.

3. We compared the *percent of latent variables annotated by at least one pathway*. For various significance threshold values that range from a p-value of $1e^{-1}$ to $1e^{-10}$, we counted the number of pathways with a p-value below the threshold for each latent variable. This

again returned a pathway coverage value for each latent variable of the embedding. We then calculated the percent of latent variables with a pathway coverage above one, which is effectively the percent of latent variables annotated by at least one pathway with a p-value below the threshold. We again repeated the calculations for each method and cancer type.

**Comparing DeepProfile pathway coverage to VAE models**

When comparing pathway enrichment of DeepProfile to VAE models, we used the gene-level attributions for each different dimensional VAE model and applied FET to obtain a p-value for each latent variable of each 600 different models. We again considered a latent variable to be significantly capturing a pathway if the FDR corrected p-value is below 0.05. DeepProfile is an ensemble model that combines 600 VAE models to define an ensemble embedding, and our aim was to show that the DeepProfile model can preserve the pathways captured by the individual VAE models. Accordingly, we used two different metrics to compare the pathway coverages of VAE models to DeepProfile:

1. We compared DeepProfile pathway coverages to the *average pathway coverages* of all 600 different VAE models. For each pathway, we calculated the percent of VAE models that captured this pathway significantly (i.e., with at least one latent variable of the embedding with an FDR corrected p-value $< 0.05$). We then compared the pathways captured by the threshold percent of the VAE models, where the threshold ranges from 50 to 90, to DeepProfile to investigate whether the pathways captured by VAE models could also be captured by DeepProfile.

2. We compared DeepProfile model to VAE models with *different dimension sizes*. For each different dimensional VAE model, e.g., a 5-dimensional VAE model, we marked a pathway to be captured if the majority of the VAE models (at least 51 of 100 models) significantly captured the pathway (FDR corrected p-value$< 0.05$). We repeated the same procedure for each of the 6 different dimensional VAE models to mark the pathways

captured by different dimensional VAE models. We then compared the pathways captured by a threshold number of different dimensional models, where the threshold ranges from 1 to 6, to the DeepProfile model in order to investigate whether the pathways captured by these different VAE models could be detected by DeepProfile as well.

**Detecting universally important genes**

To detect the highest-scoring genes across all cancer types, we used the gene-level attributions of DeepProfile latent variables. We calculated the average attribution score across all latent variables to define an overall importance score for each gene, and we converted these scores to percentile scores, where the highest scored gene takes value of 100 and the lowest scored gene takes value of 0. Once we separately obtained these percentile scores for each gene for each cancer type, we calculated the average percentile score across 18 cancer types as the universal percentile score of a gene. We could then sort the genes by their universal percentile scores to detect the top universally important ones. We generated a network of the top 100 universally important genes using STRING[21] with a medium confidence level of 0.4, using all interaction sources and eliminating disconnected latent variables. We visualized the network using Cytoscape[22].

To detect the pathways enriched for the top universally important genes, we used Fisher's Exact Test [17] *fisher_exact* method from Python's *scipy.stats* module on the top 100 universally important genes using Reactome [12], BioCarta [13], KEGG [14], and GO Biological Process (BP)[23] gene sets from version 6.2 of MSigDB [15,16]. We applied FDR correction across all pathways.

Furthermore, to detect whether DeepProfile's universally important genes are enriched for various immune cell type signatures, we collected gene signatures of T cells, B cells, neutrophils, and macrophages[24] and obtained a total of 108 genes available in our training expression dataset. We again used FET for the top 100 universally important genes using these markers to calculate enrichment score. Additionally, we incorporated the pre-computed immune cell fractions from the PanCan Immunity paper in our analysis[25]. We mapped these immune cell fractions to the TCGA gene expression data used in our study. To assess whether the top 100 DeepProfile genes could readily be explained by the genes identified as immune cell related in the PanCan study, we calculated Pearson's correlations between the fractions of four major immune cell types—T cells, B cells, neutrophils, and macrophages, as well as their subtypes—and all genes across all TCGA samples. In this analysis, the top correlated genes are likely to be specifically expressed by those immune cells. Then, for each immune cell type, we conducted FETs to assess the overlap between its top 100 correlated genes and the top 100 genes identified by our DeepProfile analysis, and applied a Bonferroni correction to adjust for multiple comparisons.

To determine whether DeepProfile's universally important genes are enriched for cell surface and cytok ine receptors, we first collected gene sets from the Cell Surface Protein Atlas (CSPA)[26], UniProt database[27], and Gene Ontology (GO). From CSPA, we downloaded the list of human surfaceome proteins and their annotations, selected the proteins with a *high confidence* CSPA category and protein probability of 1.0, and obtained a list of 555 human surface proteins. From the UniProt database, we downloaded human cell surface receptors using the keyword 'cell surface receptor,' selected the reviewed proteins, and obtained a list of 1,307 genes. Similarly, we downloaded human cytokine receptors using the keyword 'cytokine receptor,' selected the reviewed proteins, and obtained a list of 773 genes. From GO, we used the gene set 'Immune response regulating cell surface receptor signaling pathway,' which has 346 genes. Note that for each gene set, we used the genes available only in DeepProfile's training expression matrix and reported the intersecting gene counts. We again ran FET for the top 100 universally important genes using these 4 distinct gene lists to calculate enrichment scores.

To compare the top universally important genes detected by DeepProfile to the universally important genes for PCA, we ran the same analysis that we applied to DeepProfile to PCA. Using the attribution scores (the absolute valued component matrices) of each gene for the PCA model, we calculated the average attribution score across all top 150 principal components, converted the scores to percentile scores, and calculated the average across 18 cancers to define a universal importance score for each gene for PCA. Similarly, we ran FET for the top 100 universally important PCA genes using KEGG, BioCarta, Reactome pathways and GO BP gene sets. We also repeated FET enrichment tests for the same 4 receptor gene lists used for DeepProfile, again using the top 100 universally important PCA genes.

**Detecting universally important pathways**

To detect the highest-scoring pathways across all cancer types, we used the pathway-level attributions we have for each DeepProfile latent variable, which contains the p-value of enrichment for each latent variable-pathway pair. To define an overall pathway enrichment score for an embedding, we selected the maximum -log10(p-value) across all latent variables for each pathway and obtained an enrichment score for each cancer type/pathway pair. We marked a pathway as being significantly captured by a cancer type if the FDR corrected p-value is below 0.05. To determine the universally important pathways, we counted the number of cancer types that significantly captured each pathway. We also recorded the average -log10(p-value) of enrichment for each pathway by taking the mean across all cancer types that significantly captured the pathway. After obtaining the number of cancers and average enrichment score for each pathway, we sorted the pathways first by the number of cancers and then by enrichment scores to get the list of universally important pathways.

**Calculating cancer character scores for pathways**

To calculate a cancer character score for each pathway, we conducted what we called a "normal tissue analysis." First, from the GTEX portal (https://gtexportal.org/home/datasets), we downloaded RNA-Seq expression (gene TPMs) with accession number phs000424.v7.p2[28] and selected the tissues corresponding to the 18 cancer types we have (see **Supplementary File 1** for the mapping of DeepProfile and GTEX tissue types and the number of samples). We preprocessed and encoded GTEX expression profiles following the same pipeline used for TCGA RNA-Seq expression and passed the expression values to our already trained DeepProfile models to generate normal tissue embeddings.

To detect how successfully each latent variable can differentiate cancer vs. normal tissue, we trained logistic regression classifiers by passing the cancer and normal tissue DeepProfile embeddings as input and predicted cancer vs. normal tissue labels. We used the Python *sklearn* library's *LogisticRegression* with *liblinear* solver and *l2* regularization, repeated the training 500 times with different random samplings, and recorded the mean of absolute value of classifier weights from all models. We defined the cancer character score of each latent variable as the absolute valued classifier weight, denoting the importance of each DeepProfile latent variable in differentiating cancer from normal tissue, where a high cancer character score would indicate that the latent variable is quite important for differentiating the tissue type.

We then mapped these latent variable-level cancer character scores to pathways to determine the cancer-tissue specificity of each pathway for each cancer type. For each pathway, we calculated the weighted average of cancer character scores using the -log10(p-value) of enrichment scores for that pathway as the weights and obtained an average cancer character score for each pathway-cancer type pair. Note that if a pathway is not enriched for any of the latent variables, we assigned it a cancer character score of 0. To define a universal cancer character score for each pathway, we calculated the average across the cancer character scores of 18 cancers, excluding the cancers with a score of 0.

**Detecting cancer-specific genes and pathways**

To identify the genes that are high scoring specifically for a certain cancer type, we used the importance scores we calculated for each gene-cancer type pair. For each gene, we calculated the difference between the percentile score of a cancer and the maximum percentile score across all other 17 cancers. These cancer-specific difference scores allowed us to detect the top cancer-specific genes for each cancer.

To calculate the enrichment score for the PAM50 genes[29], we ran FET for the top N highest scoring genes for breast cancer where N ranges from 1 to 1000. We then applied Bonferroni correction over all thresholds to report the final p-value of enrichment.

To detect cancer-specific pathways, we used the -log10(p-value) of enrichment scores we calculated for each pathway/cancer type pair and calculated the difference between the enrichment score for one cancer type and the maximum enrichment score across all other 17 cancers. Thus, we obtained a cancer-specific difference score for each cancer type and for each pathway, allowing us to detect the pathways that are cancer type-specific. We also assigned a cancer character score to each of these cancer-specific pathways using the absolute-valued classifier weights from the normal tissue analysis after converting them to percentile scores, as well.

**Pan-cancer survival and mutation analysis**

With the goal of associating each pathway with patient survival, we used the TCGA RNA-Seq DeepProfile embeddings we learned for 18 cancers along with the survival status. We separately fitted univariate Cox regression models[30] to each DeepProfile latent variable. We used the R *survival* library's *coxph* method, recorded p-values of model coefficients for each latent variable, and applied FDR correction over all latent variables for each pathway. We repeated the model trainings for each cancer type.

To detect the pathways determining patient survival, we mapped these latent variable-level survival scores to pathways. For each pathway, we calculated the weighted average of -log10(survival p-values) across all latent variables using the -log10(enrichment p-values) as the weights (see **Extended Data Fig. 6**). When none of the latent variables could significantly capture a pathway (FDR corrected p-value < 0.05), we assigned a survival score of 0 to that pathway. We also masked the pathway enrichment p-values with a z-score below 0.25 to prevent the involvement of lowly ranked pathways in the average score calculation. We calculated the average survival -log10(p-value) for each pathway/cancer type pair. Since the survival analysis using enrichment scores from FET did not provide a rich enrichment for survival, we repeated the FET using a broader set of top genes (4 * average pathway size) in order to run an informative survival analysis. This pipeline let us define a survival p-value for each pathway and for each cancer type. We then marked a pathway to be relevant to survival if both the FDR corrected enrichment p-value and the survival p-value were below 0.05.

To detect universally survival-related pathways, for each pathway, we counted the number of cancers (of the 18 cancers) with significant survival scores. We also calculated the average enrichment -log10(p-value) and the average survival -log10(p-value) across all the cancer types detected to be significantly associated with survival. Sorting the pathways by the number of cancer types and then by the average survival score provided us with the top universally survival-associated pathways. We visualized the top 20 universally survival-associated pathways using Cytoscape[22] EnrichmentMap tool[31] with latent variable cutoff value of 1.0 and edge cutoff similarity value of 0.5, where the connections between pathways were determined by the Jaccard similarity of gene memberships of pathways.

To conduct mutation analysis, we downloaded TCGA mutation profiles for all cancer types from Broad Institute data version 2016_01_28 (https://gdac.broadinstitute.org/) generated by the TCGA Research Network (https://www.cancer.gov/tcga/) (see **Supplementary File 1** for the mapping of cancer types and the number of samples for each cancer). We selected the samples for which we have both expression measurements and tumor mutational burden (TMB) data available and calculated the total number of mutations for each cancer sample by summing the number of mutations for each gene. If k different mutations occurred for one gene, we increased the total mutation count by k.

To assign a mutation association score to each DeepProfile latent variable, we calculated the Pearson correlation, using the Python *scipy.stats* library's *pearsonr* method, between each latent variable of DeepProfile embedding and the log of total mutation count after eliminating outlier mutation scores beyond the 95% confidence level (z-score > 1.96). We repeated these experiments for each of the 18 cancer types and obtained latent variable-level TMB correlation p-values.

To map the latent variable-level p-values to pathways, we repeated the same procedure we followed for the survival analysis: we calculated the weighted average -log10(TMB p-value) across all latent variables for each pathway, where the weights were defined as -log10(enrichment p-value) of the latent variables. We again calculated the average enrichment and TMB -log10(p-values) across all cancer types with significant scores (FDR corrected p-value < 0.05) and detected the number of cancers significantly associated with TMB for each pathway. We visualized the top 20 mutation-associated pathways using Cytoscape [22] EnrichmentMap tool [31] with the same setting as the survival-associated pathways network.

**Downstream survival analysis**

For pathways detected to be relevant to patient prognosis, we conducted a downstream survival analysis independent from DeepProfile model. We first fitted univariate Cox regression models[30] to each of the 23 genes included in the KEGG mismatch repair pathway and 91 genes included in the Reactome MHC class II antigen presentation pathway using TCGA RNA-Seq profiles. We used the R *survival* library's *coxph* method to predict survival, trained the models separately for each cancer type, and recorded z-scores to get the direction of association. After obtaining survival z-scores from Cox models, we created heatmaps of gene-level survival scores for 18 cancers by clustering the genes with hierarchical clustering.

To investigate the association of average expression of the selected pathways with survival, we first calculated the average expression of genes from the KEGG mismatch repair pathway and the average expression of HLA-D genes (HLA-DMA, HLA-DMB, HLA-DOA, HLA-DOB, HLA-DPA1, HLA-DPB1, HLA-DQA1, HLA-DQA2, HLA-DRB1, HLA-DRB5) from the Reactome MHC class II antigen presentation pathway across all TCGA samples with survival record. We then created Kaplan-Meier plots[32] for each pathway using the calculated average expression values. We used the Python *lifelines* library's *KaplanMeierFitter* class for generating Kaplan-Meier survival plots. When generating the plots, we separated the patients into two groups based on their average expressions: one group with expression above mean + standard deviation and one group with expression below -(mean + standard deviation). We then fitted Kaplan-Meier models to these 2 groups. We also recorded the p-values using the *lifelines logrank_test,* which tests how significantly the two curves are separated from each other.

To detect the immune cell type responsible from expression HLA-D genes, we first calculated the average expression of each gene across all TCGA samples for each cancer. Note that we performed the mean operation prior to preprocessing the expression matrices. We sorted the genes by their average expression and converted the rankings to percentile scores. To order the importance of different immune cell types, we calculated the average gene percentile score of the gene signatures for each immune cell type, which we defined as XCR1 and CLEC9A for dendritic cells; MS4A1, CD79A, and PAX5 for B cells; and CD163, CD68, CSF1R for macrophages. To measure the association between the immune cells and HLA-D expression, we measured the Pearson correlation between average expression of the cell type signatures listed above and the average expression of HLA-D genes using the Python *scipy.stats* library's *pearsonr* method.

For the macrophage analysis, we downloaded the gene signatures for pro-inflammatory and immunosuppressive macrophages[33] and used the list of unique genes for each macrophage group (CD40, CXCL9, CXCL10, CXCL11, SLAMF1, and TNIP3 for pro-inflammatory macrophages; CFP, HRH1, NPL, PDCD1LG2, and RENBP for immunosuppressive macrophages) to again measure gene ranking percentile scores. We conducted the same analysis as we did for immune cell types: we measured the average expression for genes, ranked the genes, and calculated the average percentile scores for the genes included in pro-inflammatory or immunosuppressive macrophage signatures. We also repeated this pro-inflammatory/immunosuppressive macrophage analysis using the extensive list of pro-inflammatory or immunosuppressive macrophage signatures[34].
