## Supplementary Appendix for "A deep profile of gene expression across 18 human cancers"

### Supplementary Information

#### Contents

1. Supplementary Note 1 Evaluating Pathway Uniqueness and Redundancy in DeepProfile Latent Variables
2. Supplementary Note 2 Analyzing DeepProfile's Performance on Gaussian Noise Data
3. Supplementary Note 3 Enhanced Analysis of Cancer Subtype Differentiation Using the Metabric Dataset
4. Supplementary Note 4 Robustness Evaluation of DeepProfile Model Ensembles
5. Supplementary Note 5 Assessing the Stability of K-means Clustering
6. Supplementary Note 6 Comparative Analysis of Pathway Enrichment Using GSEA and Fisher's Exact Test
7. Supplementary Note 7 Survival Prediction Analysis Using TCGA Data
8. Supplementary Figures and Tables
9. Supplementary Files List

#### Supplementary Note 1 Evaluating Pathway Uniqueness and Redundancy in DeepProfile Latent Variables

We conducted an in-depth analysis of the pathway overlap among latent variables for the KEGG, BioCarta, and Reactome Pathways analysis. Specifically, we calculated the percentage of identified pathways that are unique to a single latent variable as well as the percentage of pathways shared by at most 10% of the latent variables (which equates to 15 latent variables for the models considered). The results are presented in **Supplementary Table 5**.

Our findings indicate that, on average, 9.01% of pathways are uniquely identified by only one latent variable. This suggests that a portion of the latent variables in DeepProfile is indeed capturing unique biological variation. Additionally, 51.70% of pathways are found by no more than 10% of latent variables, indicating a modest level of redundancy and supporting the model's ability to differentiate between different sources of biological variation.

Furthermore, to address concerns about redundancy among latent variables, we analyzed the average paired overlap ratio for the top 51 genes—the number of genes considered in Fisher's exact test—across all 150 latent variables. The results, presented in **Supplementary Table 5**, indicate that for most cancers, the top gene overlap ratio ranges from 10%-15%. This demonstrates that the majority of genes identified by the latent variables are distinct, reinforcing the model's ability to effectively disentangle complex gene expression data into meaningful and non-redundant biological insights.

#### Supplementary Note 2 Analyzing DeepProfile's Performance on Gaussian Noise Data

In an effort to rigorously test DeepProfile's capability to discern true biological signals from noise, we trained the model on datasets composed exclusively of Gaussian noise. We chose breast cancer (largest sample size), sarcoma (average sample size), and bladder cancer (smallest sample size) for this analysis. The results, detailed in **Supplementary Fig. 7**, indicate that very few pathways are identified by the latent variables of the models trained on random noise. Additionally, these latent variables rarely identified even one pathway. This outcome strongly suggests that the significant pathway enrichment observed in our original models is not a result of chance, but a consequence of the model’s capability to capture meaningful biological variation in gene expression data.

#### Supplementary Note 3 Enhanced Analysis of Cancer Subtype Differentiation Using the Metabric Dataset

In our study, we explored the abilities of DeepProfile, PCA, ICA, and RP to distinguish cancer subtypes, leveraging the Metabric dataset renowned for its detailed subtype labels in breast cancer. The incorporation of the Metabric dataset enhanced our initial analysis, providing a broader platform for a more comprehensive evaluation of the performance of these models in the context of cancer subtype identification.

Embeddings for each model, initially trained on a robust dataset, were generated and subsequently applied to the Metabric dataset. The effectiveness of these models in differentiating cancer subtypes was evaluated through both visual and quantitative analyses. t-SNE plots were utilized to visually demonstrate the capabilities of DeepProfile and PCA in subtype differentiation (**Supplementary Fig. 8A-D**), in comparison to the performances of ICA and RP. These qualitative evaluations were complemented by Silhouette Width scores (**Supplementary Fig. 8E**), a metric indicating the degree of clustering efficacy within subtypes. Notably, DeepProfile and PCA both achieved positive scores, signaling effective clustering, with DeepProfile exhibiting particularly high proficiency.

In addition, we employed XGBoost models, training them with embeddings derived from each method, to classify cancer subtypes. Our findings underscore DeepProfile's exceptional ability to identify cancer subtypes, demonstrated by its superior classification accuracy, F1 scores, average sensitivity, and specificity, as well as the lowest log loss (**Supplementary Fig. 8F-J**). While DeepProfile demonstrated better performance in identifying subtypes, PCA also exhibited commendable effectiveness in this regard, indicating its ability in cancer subtype identification.

#### Supplementary Note 4 Robustness Evaluation of DeepProfile Model Ensembles

In our evaluation of the DeepProfile model's robustness, we undertook an in-depth analysis focusing on the consistency of gene ranking across various ensembles of VAE models. The analysis was structured to compare the top-ranking genes identified by each ensemble configuration, particularly observing changes upon the inclusion of additional models. We calculated the shared genes within the top 100 rankings as more models are incorporated into the ensemble. This is done by comparing the top 100 genes identified by one ensemble against the top 100 genes identified by another ensemble after adding an additional model.

The findings from this analysis, presented in **Supplementary Fig. 9**, show that with the inclusion of more models in the ensemble, there is a notable increase in the number of overlapping genes in the top rankings. This trend continues such that when the ensemble consists of 20 models, the overlap of genes in the top 100 reaches over 95%. This substantial overlap attests to the robustness of the latent space captured by the DeepProfile VAEs, indicating that our model is consistently identifying a core set of significant genes. These results suggest that our model's output is stable across different initializations and iterations, reinforcing the validity of the genes identified as being integral to the underlying biological context.

#### Supplementary Note 5 Assessing the Stability of K-means Clustering

Given the inherent variability of k-means, due to its reliance on initial centroids, especially in smaller datasets, we evaluate the stability of k-means clustering by rerunning k-means 10 times across all cancer types and calculated the average Normalized Mutual Information (NMI) scores between the clusters from each new k-means run and the original model. As shown in **Supplementary Fig. 10A**, our results indicate that the NMI scores consistently hover around 0.5, suggesting good stability.

To further investigate, we focused on two cancer types with the lowest NMI scores, namely lung and brain cancer, and also a cancer type with the smallest sample size. Our aim was to compare the identified genes and pathways between the new k-means models and the original. As displayed in **Supplementary Fig. 10B-G**, there's a significant overlap. For the chosen cancer types, even for the top 40 genes, about 90% of them match across runs. This overlap increases to over 95% for the top 200 genes. In terms of pathways, more than 80% overlap for top 20 pathways, and this goes up to over 85% for the top 100 pathways.

These results highlight that, despite k-means' potential variability, our model consistently identifies genes and pathways across different runs, ensuring stable latent variable representations.

#### Supplementary Note 6 Comparative Analysis of Pathway Enrichment Using GSEA and Fisher's Exact Test

To enhance our pathway enrichment analysis, we utilized the widely recognized Gene Set Enrichment Analysis (GSEA) on ranked gene lists from DeepProfile, comparing the outcomes with those obtained using Fisher's Exact test. We applied GSEA to ranked gene lists from the DeepProfile model, focusing on pathways with FDR-corrected p-values below 0.05 for statistical robustness. The results, presented in **Supplementary Table 6**, show a significant overlap between the pathways identified by both GSEA and Fisher's Exact Test, averaging a concordance rate of approximately 77%.

This high degree of overlap validates our initial findings and demonstrates the consistency of biologically significant pathway identification between these two distinct methodological approaches. The correlation between GSEA and Fisher's Exact Test results not only confirms the validity of our earlier results but also strengthens our confidence in the biological relevance of the identified pathways. This comprehensive analysis supports the use of Fisher's Exact Test in our study and reinforces the integrity and reliability of our findings.

#### Supplementary Note 7 Survival Prediction Analysis Using TCGA Data

We utilized data from The Cancer Genome Atlas (TCGA) to evaluate the survival prediction ability of DeepProfile. Our analysis included cancer types with over 500 samples. The TCGA expression data was transformed into embeddings using DeepProfile models trained on GEO data. These embeddings served as inputs for Cox regression through the XGBoost model, allowing for a comparative analysis with embeddings generated by other methods. This comparison included both deep learning models (AE, DAE and VAE) and linear models (random projections, PCA, and ICA).

For methodological rigor, the dataset was divided into 81% for training, 9% for validation, and 10% for testing, with each cancer type undergoing separate training. We repeated this process across 10 different train/validation/test splits, basing our conclusions on the average C-Index. Notably, DeepProfile outperformed other methods in 4 out of the 7 cancer types tested (**Supplementary Fig. 11**), demonstrating its enhanced capability in predicting survival outcomes compared to other dimensionality reduction techniques.

### Supplementary Figures and Tables

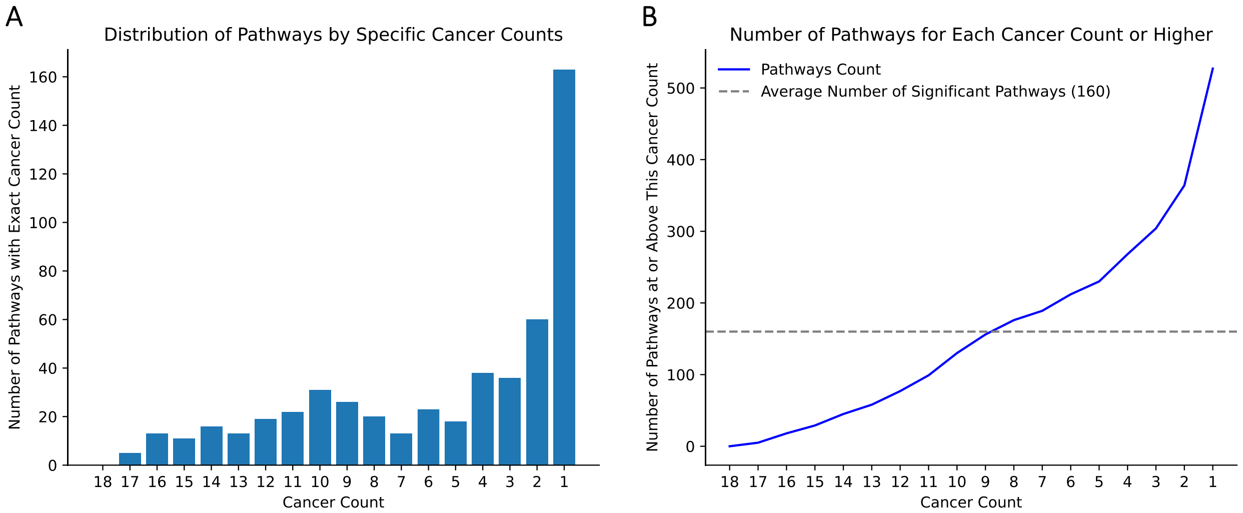


**Supplementary Fig. 1: Analysis of Pathway Distribution Across Different Cancer Counts. A** The distribution of pathways by specific cancer counts, depicting the number of pathways that are associated with exactly each cancer count from 1 to 18. **B** The number of pathways identified for each cancer count or higher, highlighting the cumulative increase. The dashed line represents the average number of significant pathways (160) identified across all cancers.

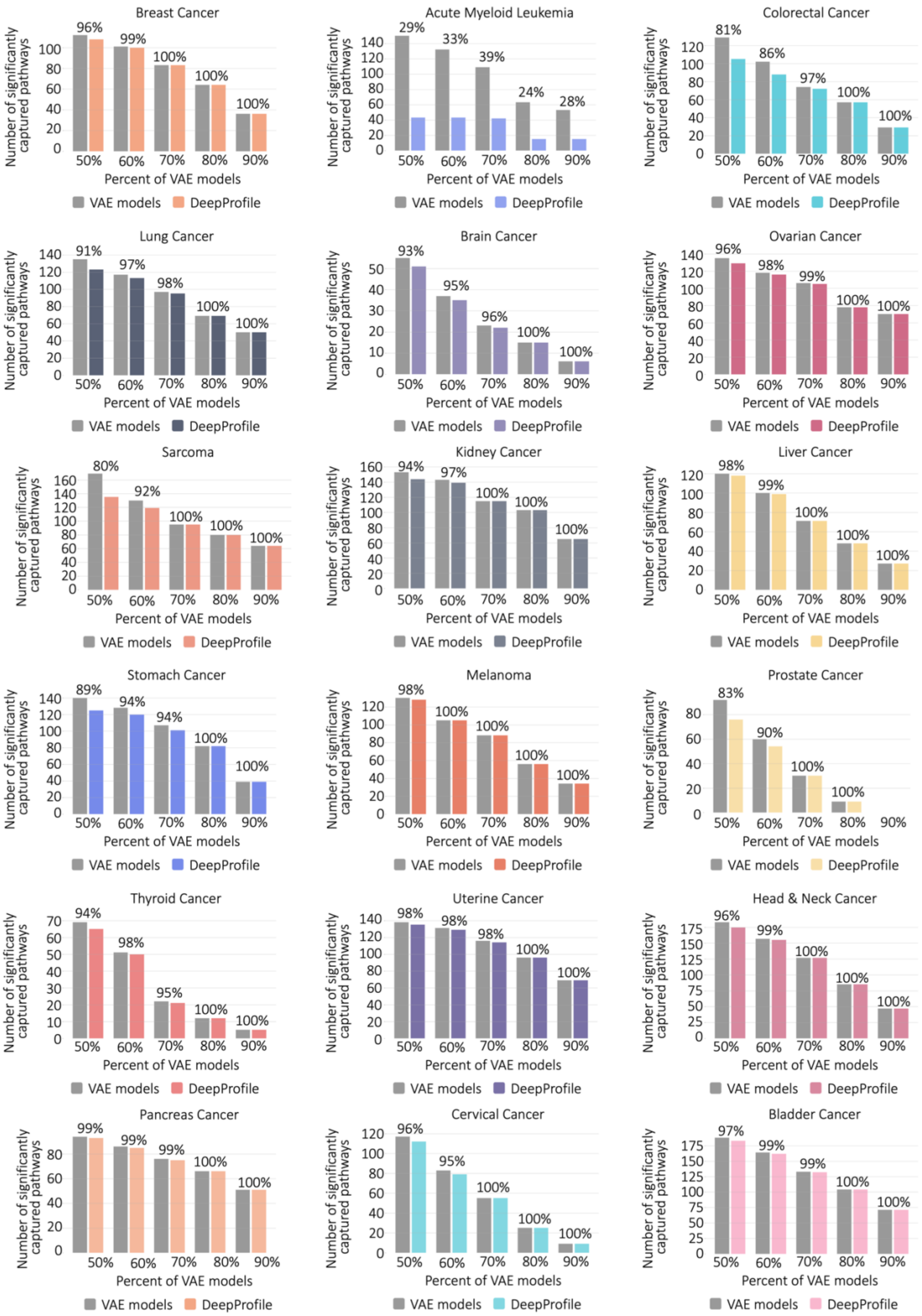

Breast Cancer
96% 99% 100% 100% 100%
Acute Myeloid Leukemia
29% 33% 39% 24% 28%
Colorectal Cancer
81% 86% 97% 100% 100%
Lung Cancer
91% 97% 98% 100% 100%
Brain Cancer
93% 95% 96% 100% 100%
Ovarian Cancer
96% 98% 99% 100% 100%
Sarcoma
80% 92% 100% 100% 100%
Kidney Cancer
94% 97% 100% 100% 100%
Liver Cancer
98% 99% 100% 100% 100%
Stomach Cancer
89% 94% 94% 100% 100%
Melanoma
98% 100% 100% 100% 100%
Prostate Cancer
83% 90% 100% 100%
Thyroid Cancer
94% 98% 95% 100% 100%
Uterine Cancer
98% 98% 98% 100% 100%
Head & Neck Cancer
96% 99% 100% 100% 100%
Pancreas Cancer
99% 99% 99% 100% 100%
Cervical Cancer
96% 95% 100% 100% 100%
Bladder Cancer
97% 99% 99% 100% 100%
Number of significantly captured pathways
50% 60% 70% 80% 90%
Percent of VAE models
VAE models
DeepProfile

**Supplementary Fig. 2: Comparison of DeepProfile's pathway coverage with individual VAE models. Related to Figure 2.** Plots of pathway coverage comparison of DeepProfile model and VAE models. For each pathway, we count the percent of all the VAE models that can significantly capture this pathway (FDR corrected p-value < 0.05). We also check if DeepProfile can significantly capture this pathway as well. For various percentage thresholds shown on the y-axis, we show the number of pathways captured by at least threshold percent of the VAE models (left bar) and the number of pathways captured by DeepProfile as well (right bar). The percentage values shown for each threshold denote the percent of pathways covered by DeepProfile. We include plots for each of the 18 cancer types. For 17 cancer types out of 18, DeepProfile can capture at least 80% of the pathways captured by at least 50% of the VAE models and 100% of the pathways captured by at least 80% of the VAE models.

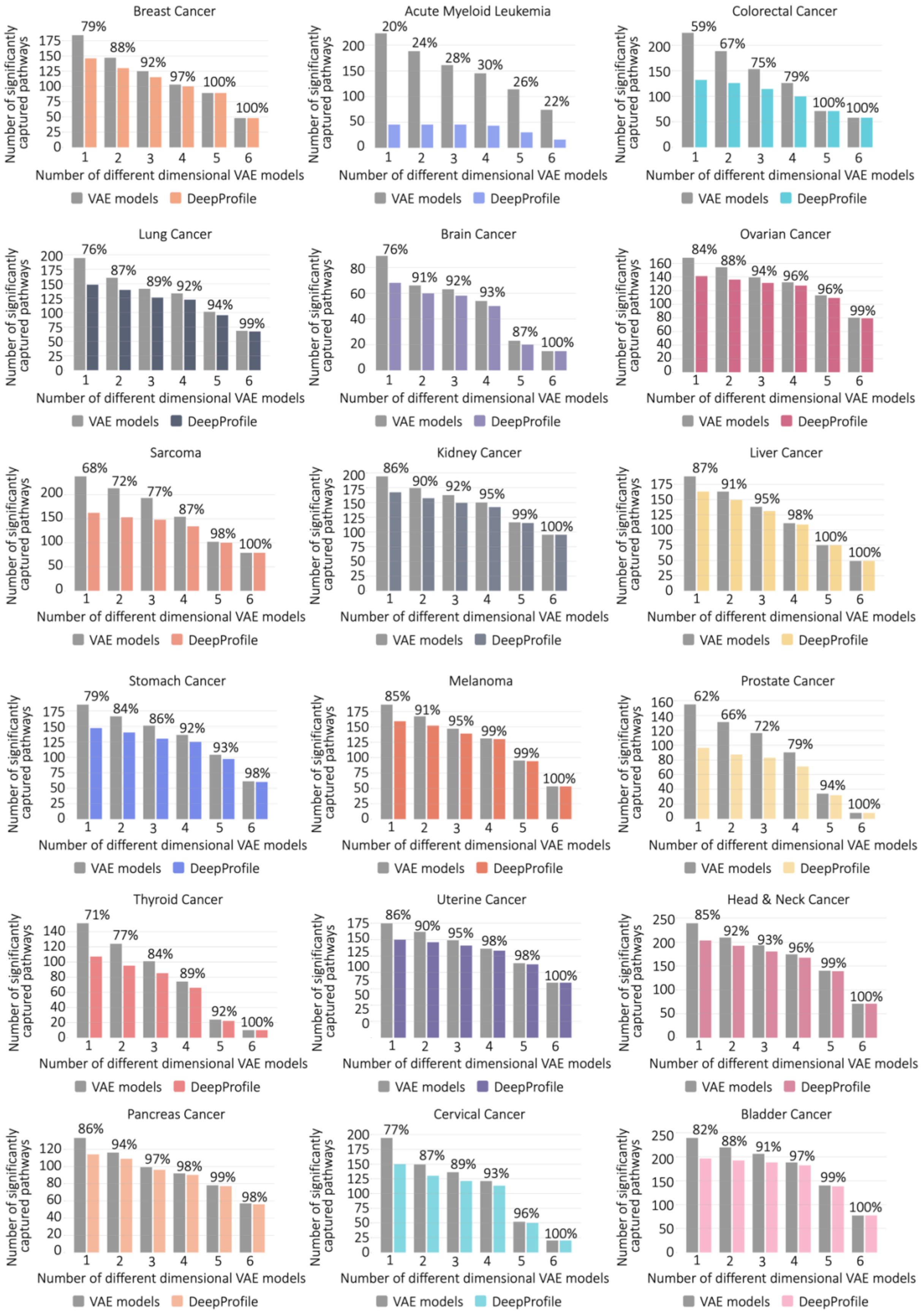
Breast Cancer
79% 88% 92% 97% 100% 100%
Acute Myeloid Leukemia
20% 24% 28% 30% 26% 22%
Colorectal Cancer
59% 67% 75% 79% 100% 100%
Lung Cancer
76% 87% 89% 92% 94% 99%
Brain Cancer
76% 91% 92% 93% 87% 100%
Ovarian Cancer
84% 88% 94% 96% 96% 99%
Sarcoma
68% 72% 77% 87% 98% 100%
Kidney Cancer
86% 90% 92% 95% 99% 100%
Liver Cancer
87% 91% 95% 98% 100% 100%
Stomach Cancer
79% 84% 86% 92% 93% 98%
Melanoma
85% 91% 95% 99% 99% 100%
Prostate Cancer
62% 66% 72% 79% 94% 100%
Thyroid Cancer
71% 77% 84% 89% 92% 100%
Uterine Cancer
86% 90% 95% 98% 98% 100%
Head & Neck Cancer
85% 92% 93% 96% 99% 100%
Pancreas Cancer
86% 94% 97% 98% 99% 98%
Cervical Cancer
77% 87% 89% 93% 96% 100%
Bladder Cancer
82% 88% 91% 97% 99% 100%
Number of significantly captured pathways
Number of different dimensional VAE models
VAE models
DeepProfile

**Supplementary Fig. 3: Comparison of DeepProfile's pathway coverage with different dimensional VAE models. Related to Figure 2.** Plots of pathway coverage comparison of DeepProfile model and different dimensional VAE models. For each pathway, we count the number of different dimensional models (out of 6 different dimension sizes) that can significantly capture this pathway (FDR corrected p-value < 0.05). We also check if DeepProfile can significantly capture this pathway as well. For various threshold counts shown on the y-axis, we show the number of pathways captured by at least threshold number of different dimensional VAE models (left bar) and the number of pathways captured by DeepProfile as well (right bar). The percentage values shown for each threshold denote the percent of pathways covered by DeepProfile. We include plots for each of the 18 cancer types. For 17 cancer types out of 18, DeepProfile can capture at least 98% of the pathways captured by all different dimensional VAE models.

| | Cell type | P-value | Bonferroni-Corrected P-value | Overlapping Genes |
|---|---|---|---|---|
| Major Immune Cell Type | B.cells | 1.00 | 1.00 | |
| | T.cells | 0.62 | 1.00 | CCL5 |
| | Macrophages | 0.62 | 1.00 | MMP14 |
| | Neutrophils | 1.00 | 1.00 | |
| Subcell Type | B.cells.naive | 1.00 | 1.00 | |
| | B.cells.memory | 1.00 | 1.00 | |
| | T.cells.CD8 | 0.62 | 1.00 | CCL5 |
| | T.cells.CD4.naive | 1.00 | 1.00 | |
| | T.cells.CD4.memory.resting | 1.00 | 1.00 | |
| | T.cells.CD4.memory.activated | 0.62 | 1.00 | CCL5 |
| | T.cells.follicular.helper | 0.62 | 1.00 | RNPS1 |
| | T.cells.regulatory..Tregs. | 0.07 | 0.91 | TAF10, RPL18, CCL5 |
| | T.cells.gamma.delta | 0.02 | 0.20 | ATP6V0E1, HINT1, SRP14, C18orf32 |
| | Macrophages.M0 | 0.62 | 1.00 | MMP14 |
| | Macrophages.M1 | 0.62 | 1.00 | CCL5 |
| | Macrophages.M2 | 1.00 | 1.00 | |
| | Neutrophils | 1.00 | 1.00 | |

**Supplementary Table 1: Results from Fisher's exact tests assessing the overlap between the top 100 genes correlated with immune cell fractions and DeepProfile's top genes.** The table displays the P values, Bonferroni-corrected P values, and the overlapping genes identified.

| | Breast | AML | Colorectal | Lung | Brain | Ovarian |
|---|---|---|---|---|---|---|
| # Principal Components | 1,000 | 1,000 | 1,000 | 1,000 | 1,000 | 1,000 |
| Total Variance Explained (%) | 77.01 | 79.38 | 81.65 | 82.56 | 84.88 | 85.85 |
| | Sarcoma | Kidney | Liver | Stomach | Melanoma | Prostate |
| # Principal Components | 1,000 | 1,000 | 1,000 | 1,000 | 1,000 | 1,000 |
| Total Variance Explained (%) | 92.75 | 89.73 | 91.03 | 93.99 | 97.69 | 98.74 |
| | Thyroid | Uterine | Head& neck | Pancreas | Cervical | Bladder |
| # Principal Components | 500 | 500 | 500 | 500 | 250 | 250 |
| Total Variance Explained (%) | 92.81 | 93.96 | 96.33 | 98.76 | 89.47 | 90.28 |

**Supplementary Table 2: Number of principal components and variance explained for each cancer type.**

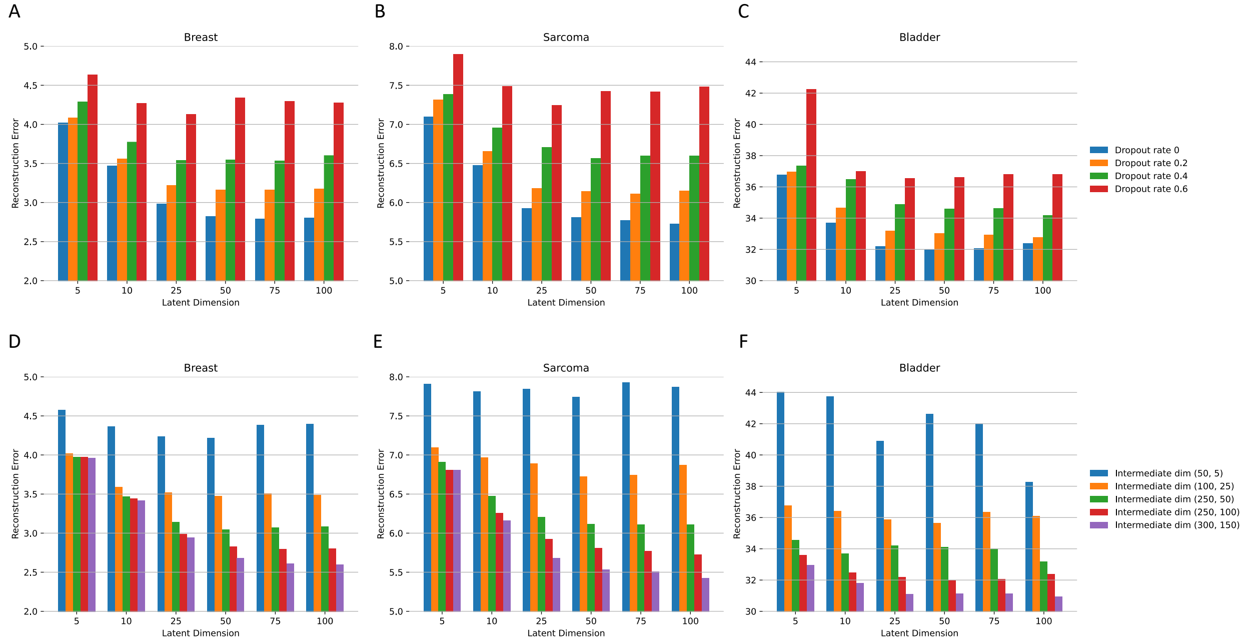


**Supplementary Fig. 4: Hyperparameter tuning for VAE models**. **A-C** Reconstruction loss (Mean Squared Error) evaluation for models with varying dropout rates, presented for breast (A), sarcoma (B), and bladder (C) cancers, which represent large, average, and small sample sizes, respectively. **D-F** Subsequent optimization of the latent dimension in the intermediate layers with a fixed dropout rate of 0.

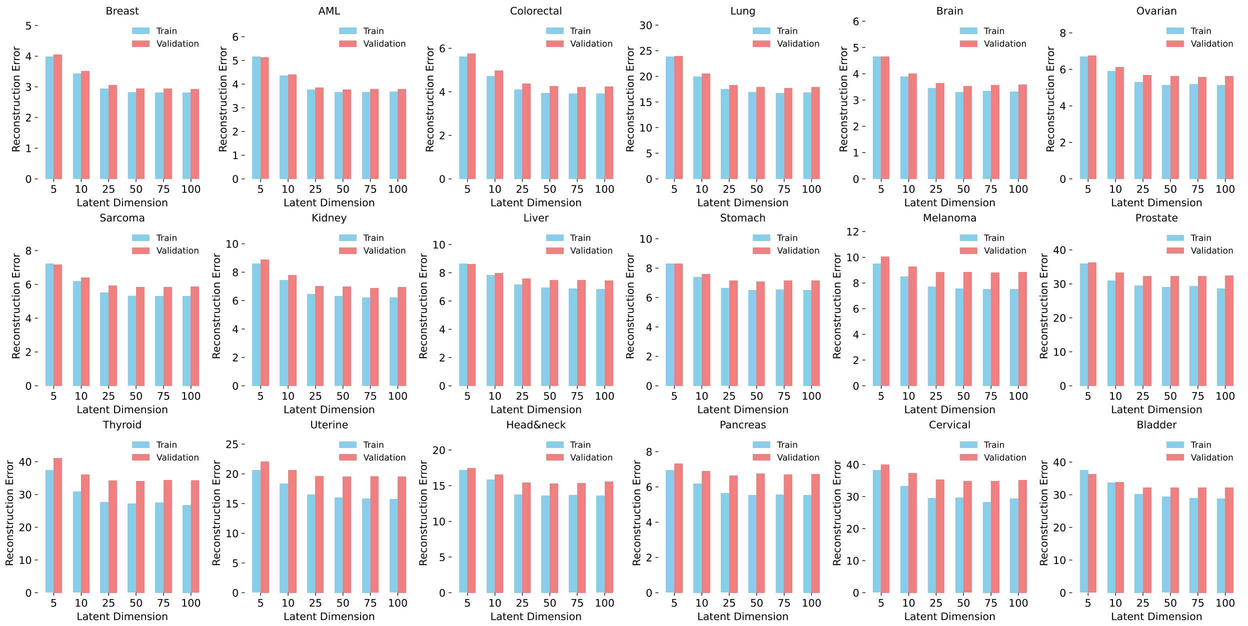


**Supplementary Fig. 5: A comparison of training and validation loss across different latent dimensions.**

| | Breast | AML | Colorectal | Lung | Brain | Ovarian |
|---|---|---|---|---|---|---|
| # Principal Components | 1,000 | 1,000 | 1,000 | 1,000 | 1,000 | 1,000 |
| # Samples | 11,963 | 6,534 | 5,616 | 4,869 | 4,282 | 2,714 |
| GPU Memory Usage (MB) | 475 | 411 | 411 | 411 | 411 | 379 |
| | Sarcoma | Kidney | Liver | Stomach | Melanoma | Prostate |
| # Principal Components | 1,000 | 1,000 | 1,000 | 1,000 | 1,000 | 1,000 |
| # Samples | 2,330 | 2,293 | 1,937 | 1,742 | 1,240 | 1,195 |
| GPU Memory Usage (MB) | 379 | 379 | 371 | 371 | 371 | 371 |
| | Thyroid | Uterine | Head& neck | Pancreas | Cervical | Bladder |
| # Principal Components | 500 | 500 | 500 | 500 | 250 | 250 |
| # Samples | 776 | 661 | 643 | 602 | 443 | 371 |
| GPU Memory Usage (MB) | 355 | 355 | 355 | 355 | 347 | 347 |

**Supplementary Table 3: GPU Memory Usage for VAE Models of DeepProfile Across Various Cancer Types.** This table details the number of principal components used, the total number of samples processed, and the GPU memory usage measured in megabytes (MB) for each cancer type, specifically for the variational autoencoder (VAE) component of the DeepProfile framework.

| | Breast | AML | Colorectal | Lung | Brain | Ovarian |
|---|---|---|---|---|---|---|
| Training time (h) | 5.36 | 2.75 | 2.37 | 2.07 | 1.83 | 1.24 |
| Testing time (h) | 0.20 | 0.12 | 0.11 | 0.11 | 0.10 | 0.08 |
| | Sarcoma | Kidney | Liver | Stomach | Melanoma | Prostate |
| Training time (h) | 1.09 | 1.07 | 1.08 | 0.99 | 0.77 | 0.74 |
| Testing time (h) | 0.08 | 0.08 | 0.07 | 0.07 | 0.07 | 0.07 |
| | Thyroid | Uterine | Head&neck | Pancreas | Cervical | Bladder |
| Training time (h) | 0.59 | 0.57 | 0.55 | 0.57 | 0.43 | 0.43 |
| Testing time (h) | 0.12 | 0.09 | 0.11 | 0.11 | 0.15 | 0.07 |

**Supplementary Table 4: Cumulative Training and Testing Times for DeepProfile Models for Each Cancer Type.** This table details the total training and testing times required for all models within the DeepProfile framework for each specific cancer type, measured in hours.

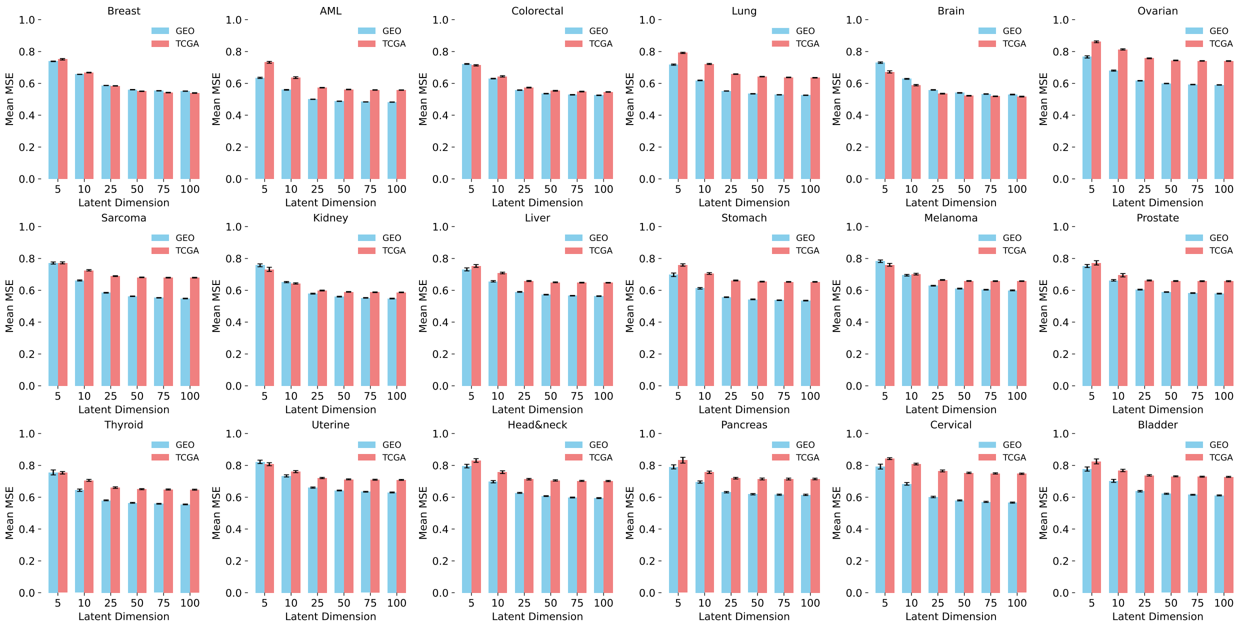


**Supplementary Fig. 6: A comparison of average Mean Square Error (MSE) between the training data (GEO) and validation data (TCGA) across different cancer types and latent dimensions.** We compare the Mean Square Error (MSE) between the GEO data, which was used for training, and the TCGA data, which was utilized for validation, on the original RNA-seq data scale. This involved transforming the reconstructed PCA data back to the original RNA-seq space before calculating the MSE. The MSE values for TCGA data are slightly higher than or comparable to those for the GEO data, showing the model's good performance on new, unseen data.

| | Breast | AML | Colorectal | Lung | Brain | Ovarian |
|---|---|---|---|---|---|---|
| 1 LV Unique Pathways | 5.10% | 6% | 17.42% | 9.04% | 12.37% | 1.28% |
| ≤ 15 LV Shared Pathways | 42.04% | 18% | 71.61% | 42.77% | 77.32% | 31.41% |
| Top Genes Overlap Ratio | 14.69% | 63.89% | 10.32% | 13.59% | 12.78% | 15.56% |
| | Sarcoma | Kidney | Liver | Stomach | Melanoma | Prostate |
| 1 LV Unique Pathways | 6.99% | 5.67% | 20.47% | 8.43% | 6.59% | 21.29% |
| ≤ 15 LV Shared Pathways | 47.85% | 43.81% | 63.72% | 48.19% | 51.10% | 74.19% |
| Top Genes Overlap Ratio | 15.32% | 11.99% | 10.44% | 11.90% | 13.34% | 12.05% |
| | Thyroid | Uterine | Head&neck | Pancreas | Cervical | Bladder |
| 1 LV Unique Pathways | 14.93% | 6.63% | 3.60% | 3.36% | 6.06% | 6.93% |
| ≤ 15 LV Shared Pathways | 82.09% | 37.35% | 58.11% | 34.45% | 67.88% | 38.61% |
| Top Genes Overlap Ratio | 9.62% | 15.17% | 16.39% | 15.27% | 13.88% | 17.02% |

**Supplementary Table 5: Evaluating Pathway Uniqueness and Redundancy in DeepProfile Latent Variables.** The percentages of pathways identified exclusively by a single latent variable (1 LV Unique Pathways %), the percentages of pathways shared by a subset of latent variables (≤ 15 LV Shared Pathways %), corresponding to 10% of the latent variables or 15 latent variables, and the average paired overlap ratio for the top 51 genes—the number of genes considered in Fisher's exact test— across all 150 latent variables (Top Genes Overlap Ratio %) for the KEGG, BioCarta, and Reactome Pathways analysis.

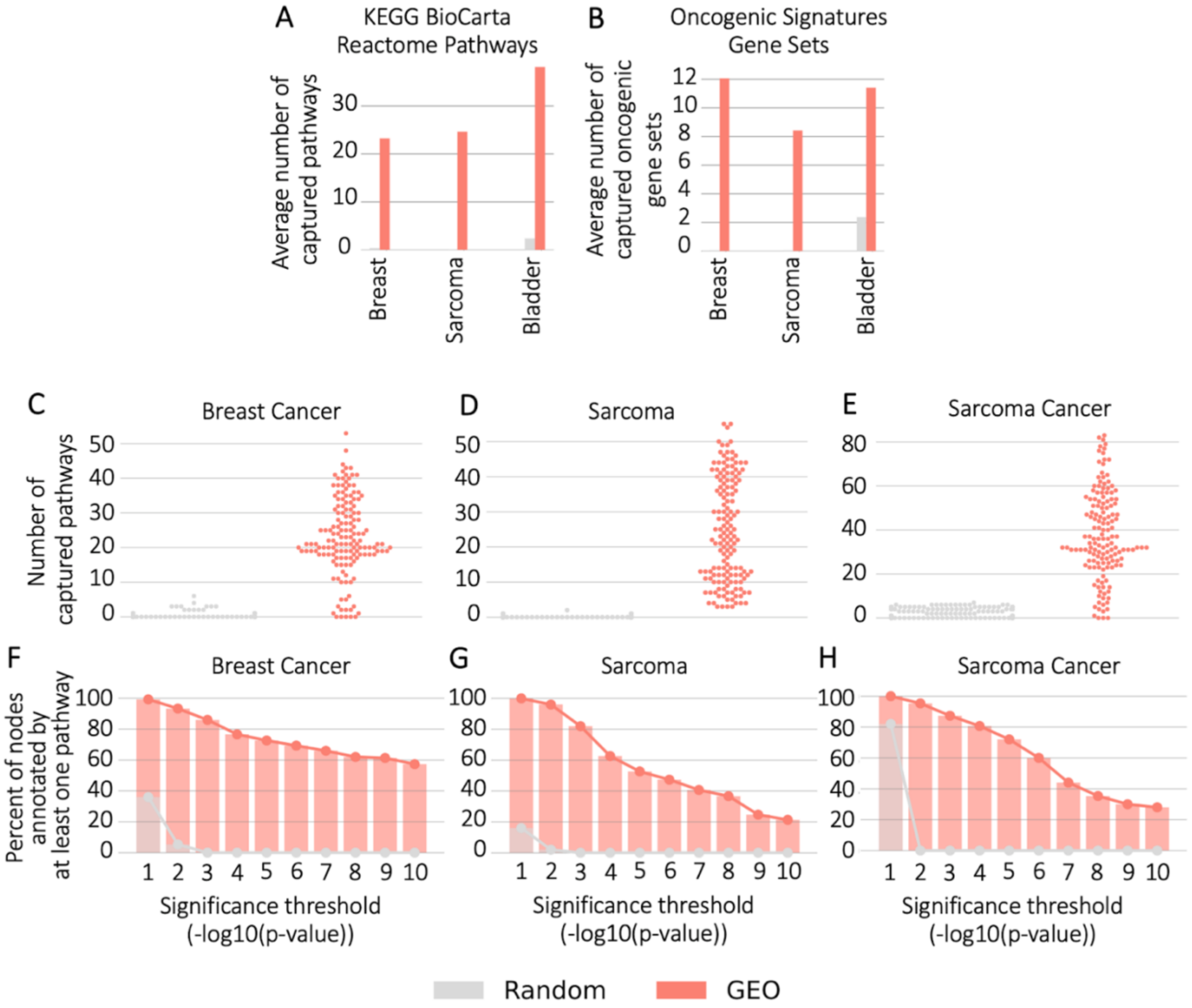


**Supplementary Fig. 7: Pathway Enrichment Analysis in DeepProfile Models Using Gaussian Noise and GEO Data. A-B** The average number of pathways with significant enrichment (FDR-corrected p-value < 0.05) identified by each latent variable in DeepProfile models trained on both Gaussian noise and GEO data. The analysis covers three cancer types: breast cancer (with the largest sample size), sarcoma (representing an average sample size), and bladder cancer (with the smallest sample size). Each latent variable of each embedding is associated with each pathway with a p-value and we count the number of pathways significantly captured by each latent variable. We then average these pathway counts over all latent variables to define the average number of pathways significantly captured by a method. **C-E** Distribution plots of number of KEGG, BioCarta, Reactome pathways significantly captured (FDR corrected p-value < 0.05) by each latent variable shown for 3 cancer types. **F-H** Comparison of the percent of latent variables annotated by at least one pathway above the significance threshold. The percent of annotated latent variables are shown for multiple significance thresholds for DeepProfile and alternative dimensionality reduction methods.

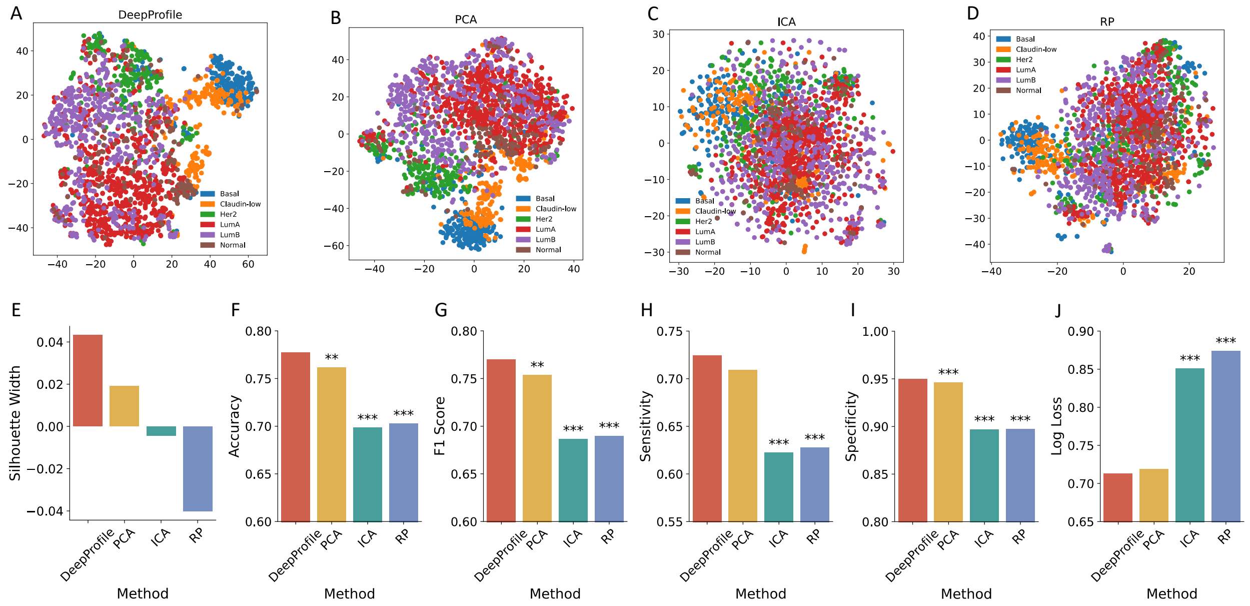


**Supplementary Fig. 8: Comparative Analysis of Gene Expression Embedding Methods for Breast Cancer Subtype Differentiation. A-D** t-SNE visualization of cancer subtype clustering by different methods: DeepProfile (A), PCA (B), ICA (C), and RP (D). Each point represents an embedded sample, colored by known breast cancer subtypes. **E** Silhouette Width comparison across methods, assessing the clustering quality of cancer subtypes in the Metabric dataset. **F-J** Accuracy, F1 score, average sensitivity, average specificity, and log loss of XGBoost models trained with DeepProfile, PCA, ICA, and RP embeddings for subtype classification, assessed across 10 train/test splits. Paired t-tests compare DeepProfile against each linear method, with '**' indicating $p<0.01$ and '***' $p<0.001$, showing statistical performance differences.

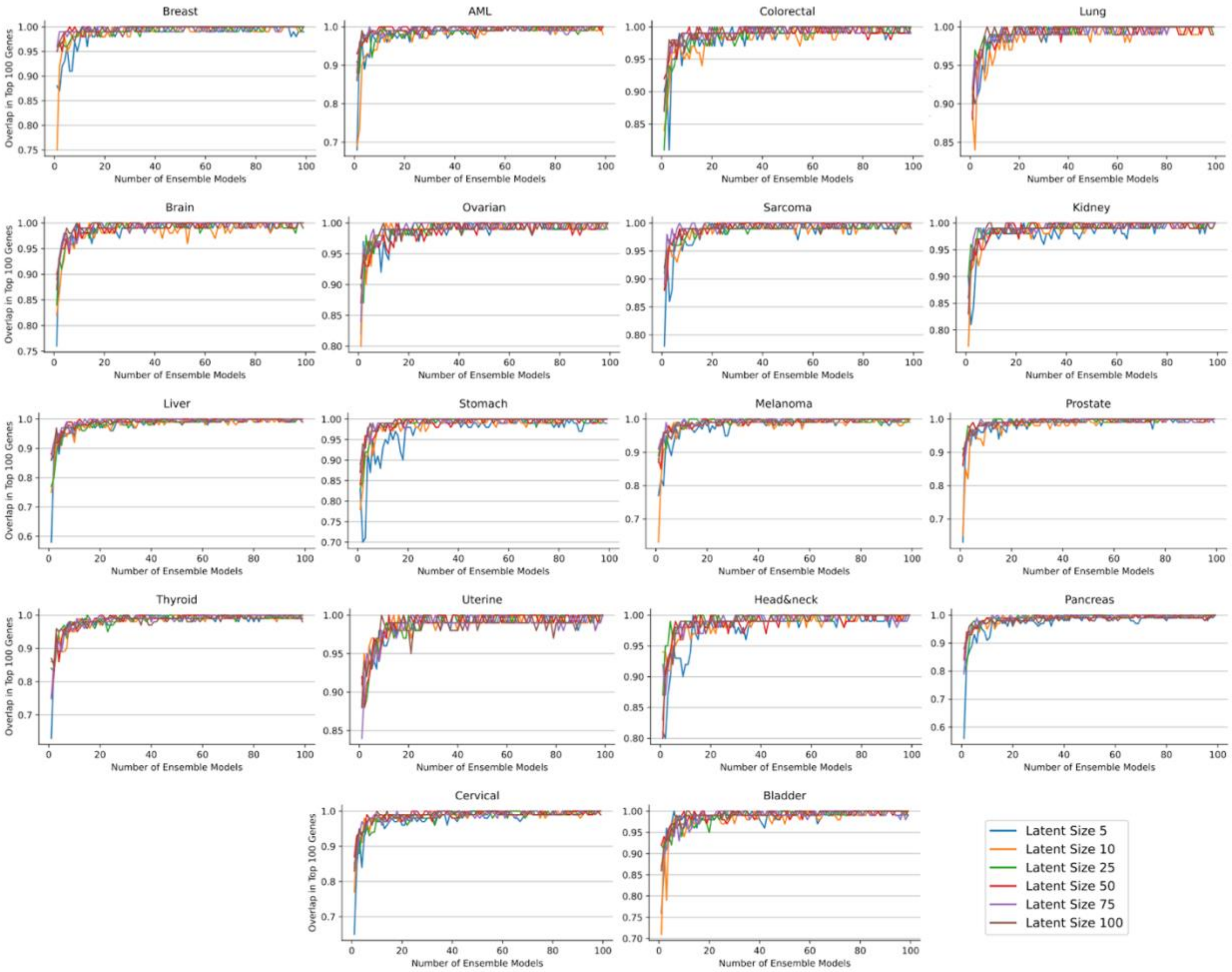


**Supplementary Fig. 9: Consistency of Gene Rankings in Ensembled VAE Models.** This figure illustrates the increasing overlap in the top 100 gene rankings as additional VAE models are ensembled. Each subplot corresponds to a different type of cancer and shows the consistency in gene identification across increasing numbers of ensembled models. The lines represent different latent space sizes (5, 10, 25, 50, 75, 100), demonstrating that larger latent spaces tend to stabilize at a high overlap percentage more rapidly. This pattern evidences the robustness of the VAE models in capturing a consistent set of significant genes across multiple training iterations and initializations.

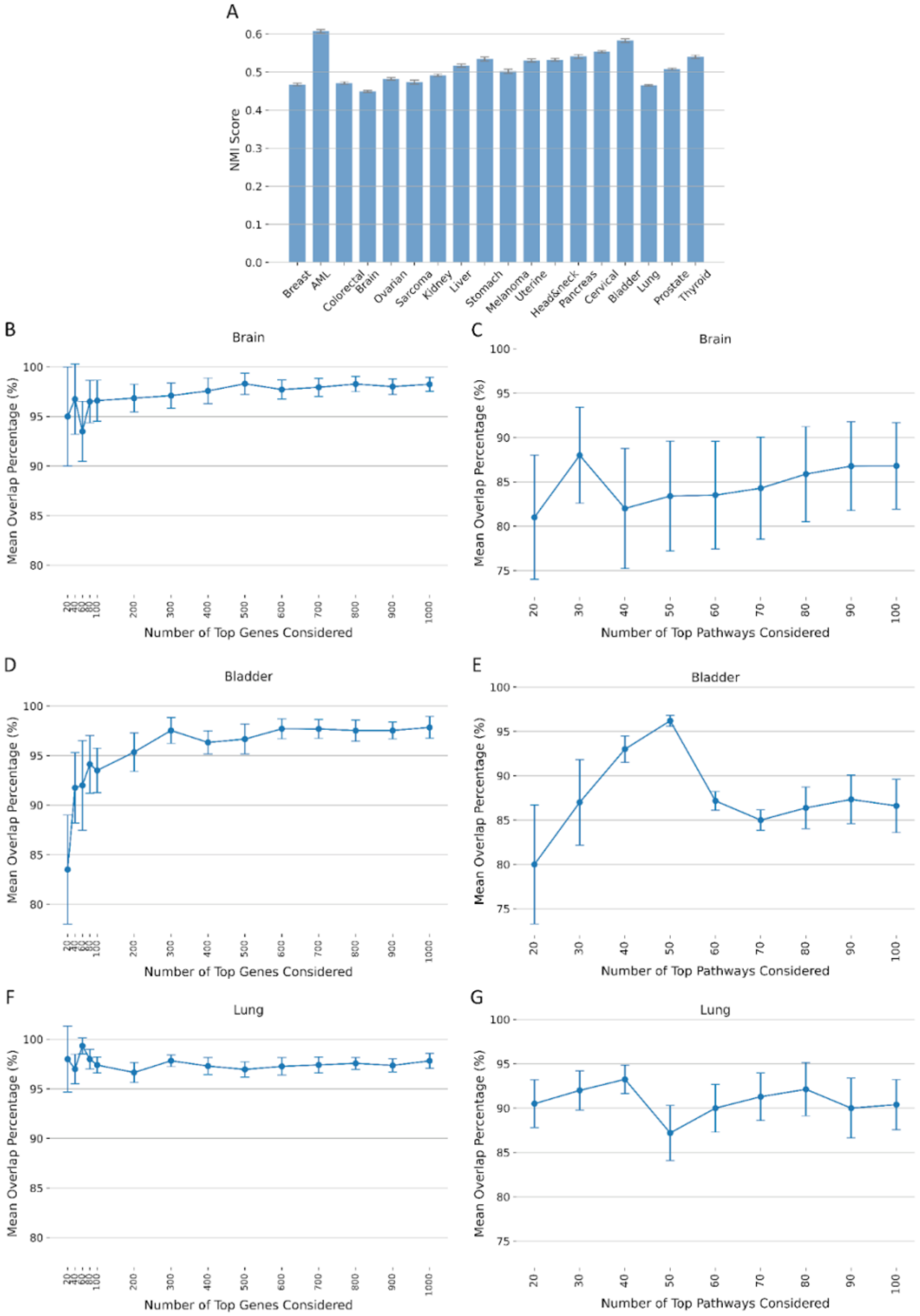


**Supplementary Fig. 10: Stability Analysis of K-means Clustering. A** Normalized Mutual Information (NMI) scores comparing the original model clusters with those from 10 new k-means runs across various cancer types. **B, D, F** Gene identification overlap in the new k-means models compared to the original. **C, E, G** Pathway identification overlap in the new k-means models compared to the original.

| | | | | | | |
|---|---|---|---|---|---|---|
| | Breast | AML | Colorectal | Lung | Brain | Ovarian |
| Percentage of Fisher's Pathways in GSEA Results | 71.97% | 66% | 72.9% | 74.10% | 72.16% | 70.51% |
| | Sarcoma | Kidney | Liver | Stomach | Melanoma | Prostate |
| Percentage of Fisher's Pathways in GSEA Results | 70.43% | 82.47% | 85.58% | 78.92% | 85.71% | 80.65% |
| | Thyroid | Uterine | Head& neck | Pancreas | Cervical | Bladder |
| Percentage of Fisher's Pathways in GSEA Results | 79.10% | 85.54% | 77.02% | 73.11% | 81.82% | 84.65% |

**Supplementary Table 6**: **Percentage of Fisher's Pathways in GSEA Results for Various Cancer Types.** This table showcases the extent of overlap between pathways identified by Fisher's Exact Test and those revealed through GSEA. It highlights the percentage of Fisher's Exact Test pathways that are also detected in GSEA results, all evaluated under the stringent statistical threshold of FDR-corrected p-values below 0.05.

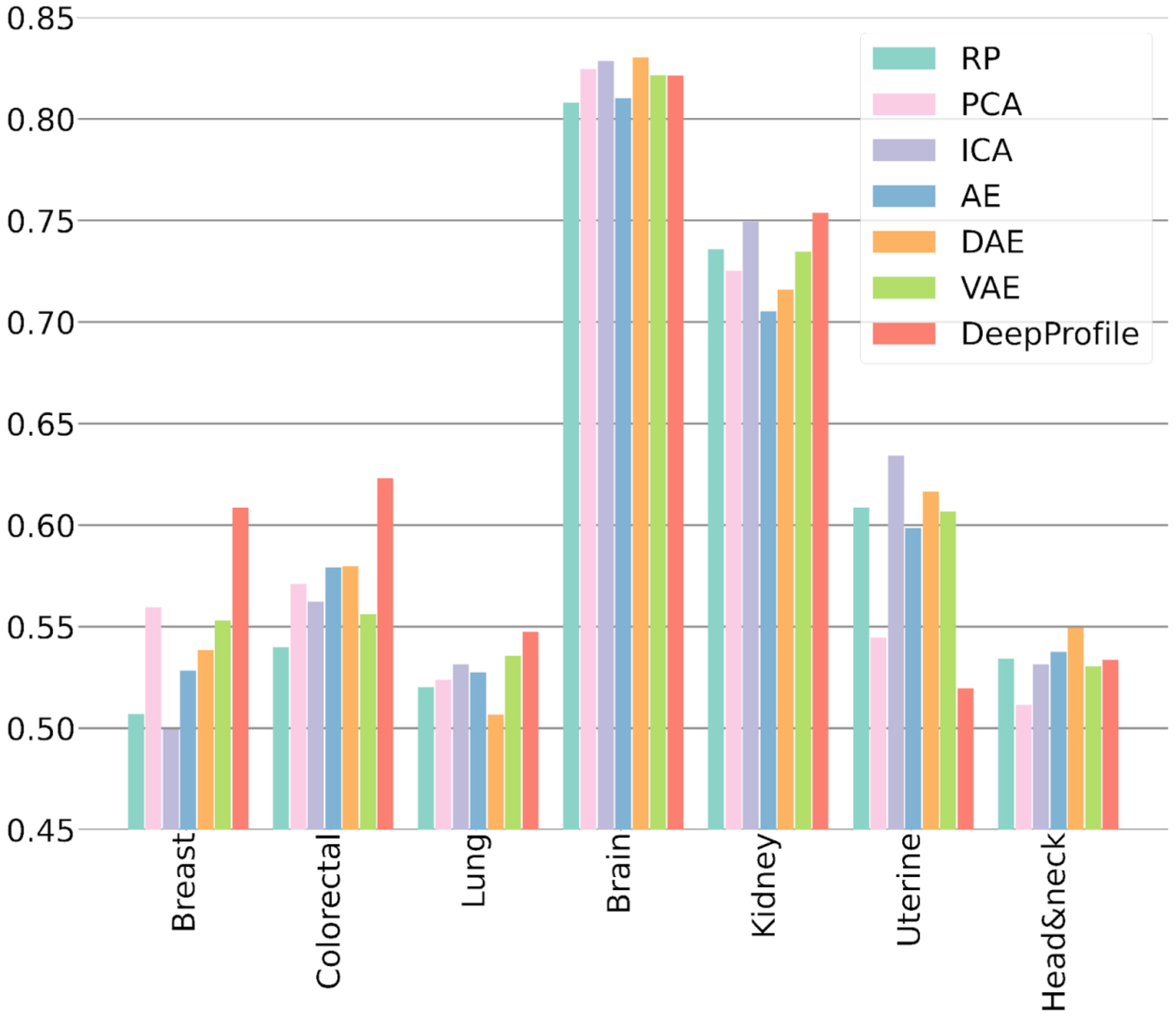


**Supplementary Fig. 11: Comparative C-index scores across various dimensionality reduction models for each cancer type on TCGA data.** The graph illustrates the average C-index of each model in predicting survival using XGBoost Cox regression. Details could be found in **Supplementary Note 7**.

#### Supplementary Files List

**Supplementary File 1 – List of GEO Series and Mappings of TCGA and GTEX for Each Cancer Type**

**Supplementary File 2 – List of Universal Gene Percentile Scores and Pathway Enrichments scores for DeepProfile and PCA**

**Supplementary File 3 – Gene Percentile Scores and Pathway Enrichment Scores for 18 Cancer Types**

**Supplementary File 4 – List of Immune Cell Marker Signatures**

**Supplementary File 5 – List of Cell Surface Markers from CSPA**

**Supplementary File 6 – Cancer-specific Genes Analysis Results**

**Supplementary File 7 – Cancer-specific Pathways Analysis Results**

**Supplementary File 8 – Cancer-specific Survival and Mutation Analysis Results**

**Supplementary File 9 – Pathway Survival and Mutation Scores**

**Supplementary File 10 – Correlations and Z-scores for Survival and Mutation**

**Supplementary File 11 – Pathway-level Survival and Mutation Analysis for 18 Cancer Types**

**Supplementary File 12 – Survival Z-scores for KEGG Mismatch Repair Genes and Reactome Mhc Class II Antigen Presentation Pathway Genes**
